# Specific and conserved patterns of microbiota-structuring by maize benzoxazinoids in the field

**DOI:** 10.1101/2020.05.03.075135

**Authors:** Selma Cadot, Hang Guan, Moritz Bigalke, Jean-Claude Walser, Georg Jander, Matthias Erb, Marcel van der Heijden, Klaus Schlaeppi

**Affiliations:** Division of Agroecology and Environment, Agroscope, Zurich, Switzerland; Institute of Geography, University of Bern, Switzerland; Genetic Diversity Centre, D-USYS, ETH Zurich, Zurich, Switzerland; Boyce Thompson Institute, Ithaca, NY, USA; Institute of Plant Sciences, University of Bern, Switzerland; Institute of Plant and Microbial Biology, University of Zurich, 8008 Zurich, Switzerland; Plant-Microbe Interactions, Institute of Environmental Biology, Faculty of Science, Utrecht University, Utrecht, The Netherlands

**Keywords:** *Zea mays*, root exudates, benzoxazinoids, rhizosphere, root microbiota

## Abstract

**Background:** Plants influence their root and rhizosphere microbial communities through the secretion of root exudates. However, how specific classes of root exudate compounds impact the assembly of these root-associated microbiotas is not well understood. Maize roots secrete benzoxazinoids (BXs), a class of indole-derived defense compounds, and thereby impact the assembly of their microbiota. Here, we investigated the broader impacts of BX exudation on root and rhizosphere microbiotas of adult maize plants grown under natural conditions at different field locations in Europe and the US. We examined the microbiotas of BX-producing and multiple BX-defective lines in two genetic backgrounds across three soil types.

**Results:** Our analysis showed that the secretion of BXs affected community composition of rhizosphere and root microbiota, with the most pronounced effects observed for root fungi. The impact of the two genetic backgrounds was weaker than that of the presence or absence of BXs, suggesting that BX exudation is a key trait by which maize structures its associated microbiota. BX-producing plants were not consistently enriching microbial lineages across the three soil types. Instead, BX exudation consistently depleted *Flavobacteriaceae* and *Comamonadaceae*, and enriched various plant pathogenic fungi in the roots.

**Conclusions:** These findings reveal that BXs have a selective impact on root and rhizosphere microbiota composition across different field locations. Taken together, this study identifies the BX pathway as an interesting breeding target to manipulate plant-microbiome interactions.

## BACKGROUND

Plants accommodate a specific and species-rich microbiota, including bacteria and fungi, on and in their roots, and in the rhizosphere. The rhizosphere refers to the soil zone surrounding the roots that is impacted by plant exudates (see below). Previous studies have shown characteristic microbiotas of root-associated compartments for plant species, including *Arabidopsis thaliana* [1, 2], *Oryza sativa* [3]*, Agave spp.* [4], *Populus deltoides* [5] and *Zea mays* [6]. Similar to the microbial communities in human or animal guts, these microbes collectively function as a microbiome and impact host performance [7]. Beneficial traits of the plant root-associated microbes are manifold and include hormone-mediated plant growth promotion [8], contribution to nutrient uptake [9–11], direct protection against pathogens [12], indirect modulation of the plant immune system to enhance pathogen resistance [13], and improving abiotic stress tolerance [14, 15]. The composition of plant root and rhizosphere microbiotas primarily reflects their origin from the surrounding soil microbiota, which is edaphically determined by the physico-chemical characteristics of a soil type [7, 16].

In addition to ‘soil type’ as a main driver of plant microbiota composition, the plant genotype has a smaller impact, explaining around 5% of the variation in microbiota composition [7, 16]. Plants mainly influence their root and rhizosphere microbial communities through the secretion of root exudates, which probably present the mechanistic link between host genetic variation and observed differences in microbiota composition between different genotypes [17]. It is likely that the combination of root exudates, the microbes’ substrate preferences and their competitiveness for the diverse carbon sources result in the specific composition of a microbiota [18]. Root exudates consist of a diverse array of exuded chemicals, with general compounds such as organic acids or sugars that mainly serve as nutritional carbon sources, and specialized compounds with semiochemical or toxic properties that further sculpt microbiota composition [17, 19]. While the plant genotype, which defines a plant’s exudation profile, is recognized as an important factor for plant microbiota assembly, relatively little is known about specific classes of exudate compounds impacting root and rhizosphere microbiota composition. It is unknown whether there are conserved microbiota responses to root exudation patterns across different soil types. The existence of widespread and conserved response patterns presents an important basis to deploy exudate chemistry for managing microbial communities.

Recent work focusing on specialized exudate compounds, for example coumarins or benzoxazinoids (BXs), points to a selective function of such secondary metabolites in plant microbiota assembly. Coumarins are abundant phenylalanine-derived specialized metabolites occurring in many plant families [20]. Recently, coumarins were reported to shape the root-associated microbiota of Arabidopsis when secreted from roots [21, 22]. Scopoletin, the predominant coumarin, excerpts direct antimicrobial activity against soil-borne fungal pathogens but not against growth-promoting and systemic resistance-inducing rhizobacteria, suggesting that plants assemble a health-promoting root microbiota through coumarin exudation [21]. Benzoxazinoids (BXs) are indole-derived specialized compounds of the *Poaceae* including major crops like maize, wheat, and rye [23]. Maize secretes substantial amounts of BXs to the rhizosphere, thereby impacting the assembly of the root and rhizosphere microbiota [24–26]. BXs are primarily known for their chemical defense functions against herbivores, pathogens, and competing plant species [27]. However, their ecological repertoire is broader [28] as they can also function as defense signaling molecules [29, 30], and phytosiderophores for iron uptake [31].

Similar to coumarins, the secretion of BXs appears to assemble more health-promoting root and rhizosphere microbiotas. We demonstrated recently that root and rhizosphere microbial communities, which were conditioned by exuded BXs, enhanced jasmonate signaling and defenses against insect herbivores in the next plant generation [24]. While these BX-driven plant-soil feedbacks present an indirect health promoting function of root and rhizosphere microbiotas, BXs also appear to function directly against soil-borne fungal pathogens. Evidence comes from recent studies comparing BX effects on root and rhizosphere microbiotas that found reduced abundances of fungal sequences with taxonomic links to plant pathogens on BX-producing plants [25, 26]. BXs were found to reduce the virulence of the plant pathogen *Agrobacterium tumefaciens* through bacteriostatic effects [32], and attracting resistance-inducing rhizobacterium *Pseudomonas putida* [33, 34]. Taken together, these findings suggest that BXs in maize root exudates function in assembling a more health-promoting root microbiota.

Several genetically different maize lines are available to study the effects of BX exudation on microbiota composition. The maize backgrounds B73 and W22 primarily accumulate the BXs 2,4-dihydroxy-7-methoxy-1,4-benzoxazin-3-one glucose (DIMBOA-Glc), DIMBOA and *N*-*O*-methylated DIMBOA-Glc (HDMBOA-Glc; [31]). Mutant lines in BX1, the first enzyme of the BX biosynthesis pathway, which converts indole-3-glycerolphosphate to indole, were identified in both backgrounds. *Bx1*(B73) and *bx1*(W22) are both deficient in the accumulation and secretion of BXs [24, 31]. BX2 encodes the second enzyme of the BX biosynthesis that converts indole to indolin-2-one. The knock-out mutant *bx2*(W22) phenocopies the BX deficiency of *bx1* mutants [31]. BX6 is a downstream enzyme in the BX pathway that acts in the multi-step conversion of DIBOA-Glc to DIMBOA-Glc. Consequently, the BX profile of the mutant *bx6*(W22) differs in its speciation from wild-type (WT) plants [31]. Overall, the mutant *bx6*(W22) produces 15% less BXs with in particular lower levels of DIMBOA and its glucoside and instead accumulates higher amounts of DIBOA-Glc (the precursor of DIMBOA-Glc).

Earlier studies focused on BX-dependent microbial feedbacks [24], were investigating 17-day-young plants [25] or were conducted in semi-artificial rhizobox systems [26] but were not comparing BX impacts on microbiotas in different soil types under field conditions. As microbiota assembly can be highly context- and soil-type dependent, we investigated the broader impacts of BXs exudation on root and rhizosphere microbiotas of 3-month-old maize plants grown under agriculturally relevant conditions at different field locations in Europe and the US. We were interested in the following specific research questions: i) How do soils, rhizosphere and roots compare in their microbiota responses to BX exudation?, ii) What is the influence of the plant genetic background on BX-mediated microbiota effects?, iii) How do different mutations in the BX biosynthesis pathway shape rhizosphere and root microbial communities?, and iv) Is there a core of microbial taxa that consistently responds to BX exudation across the different soils?

We approached these questions by conducting two field experiments, one in a loamy soil in Aurora, NY (USA) and the other in a clay loam soil in Reckenholz near Zurich (Switzerland), where we grew the various mutant lines (*bx1*, *bx2*, and *bx6*) in the two genetic backgrounds, B73 and W22. We profiled soil, rhizosphere and root microbial communities utilizing the same method as in our earlier work and compared the data from the two new experiments to the existing microbiota profiles from plants grown in a field in Changins, Switzerland (clay loam soil; [24]). In summary, we show that the BXs affected community composition of rhizosphere and root compartments and we found that BX-producing plants accumulate lower levels of *Flavobacteriaceae*, *Comamonadaceae,* and an enriched plant pathogenic fungi as a conserved microbiota response pattern.

## METHODS

### Plant genotypes

We examined the *Zea mays* L. mutant line *bx1*(B73), representing a near-isogenic line that was backcrossed five times to its wild-type (WT) background B73 [35]. We also worked with the mutants *bx1*(W22), *bx2*(W22) and *bx6*(W22) and their WT background W22 [36, 37]. The mutants in W22 have slightly different genetic backgrounds, because they were identified in *Ds* transposon lines with different anthocyanin genes (a1 and r1) as transposon launch sites. The mutants *bx1*(W22) and *bx6*(W22) were identified in W22 r1-sc:m3 (also known as T43) whereas *bx2*(W22) originates from W22 a1-m3. We utilized the W22 line with the r1-sc:m3 mutation as reference.

### Field experiments

The experiment in Reckenholz (Switzerland) was conducted in 2016 and consisted of WT and *bx1* mutant lines in both genetic backgrounds (B73 and W22), which we grew as single plants separated by maize hybrids in a field at Agroscope (Parcel 209, 47°25’34.5”N 8°31’05.9”E). Seed stock leftovers of various maize hybrid varieties were mixed and planted as buffer between the test plants. To avoid having the maize hybrids outgrow the B73 and W22 inbred lines, we pre-grew these plants for three weeks in the greenhouse in 7 x 7 x 9 cm pots, which were filled with 5-mm-sieved field soil from Parcel 209. The field was ploughed (20 cm depth) and harrowed on April 20^th^ and then on May 9^th^ after harrowing a second time, the seeds of the hybrid plants were sown. On May 27^th^, we replaced individual hybrid plants by transplanting the pre-grown inbred line plants (including the soil in their pots) to the field. The field setup was such that we grew 4 rows containing test plants, which were separated by one buffer row of hybrid plants between them. Two rows of hybrid plants surrounded the field site (**Fig. S1A**). The four genotypes B73, *bx1*(B73), W22, *bx1*(W22) were mixed and spaced within the rows by 1 m with hybrid plants in between them. Field management followed conventional farming practices consisting of herbicide applications (June 27^th^: 1.5 l/ha of Landis (44 g/l Tembotrione and 22 g/l Isoxadifen-ethyl) and 1.5 l/ha Terbuthylazin (333 g/l) and Flufenacet (200 g/l) and mineral fertilizer application, 2.5 dt/ha of magnesium-ammonium nitrate and 1 dt/ha of urea (June 24^th^), and 1 dt/ha of urea (July 6^th^). Preceding crops were winter barley (2015) and potato (2014). Single plants were harvested on August 08^th^, three months after sowing, before they started flowering.

The maize lines W22, *bx1*(W22), *bx2*(W22) and *bx6*(W22) were sampled in a field experiment in Aurora (United States, 2016) at the Cornell Musgrave Research Farm in Aurora, NY (Field U, 42.43’ 23’’ 84°N; −76.39’ 28’’ 84°E). The field was chiseled, ploughed to 18 cm, and disked for seedbed preparation. Sowing was on May 18^th^. The four genotypes were planted together with many other maize lines for seed bulking and all maize lines were grown in rows of 5.6 m length (∼20 plants per row) per block (**Fig. S1B**). We collected the mutants *bx1(W22), bx2*(W22) and *bx6*(W22) from two different blocks. Field management was according to conventional farming practices consisting of weed control (herbicide application of 0.6 l/ha Metolachlor and 0.9 l/ha Atrazine), and fertilization with 0.9 dt/ha 10-20-20 NPK at planting and 0.54 dt/ha N in the form of urea and ammonium nitrate a month after planting. The field was rotated with maize and soybean in the past and had maize (2015) as last pre-crop before the experiment. The plants were at flowering stage when single plants were harvested on September 16^th^, 4 months after sowing.

The setup and field management of the experiment in Changins (Switzerland) was described earlier [24].

### Soil analysis

We determined the soil characteristics of the three fields at the Labor für Boden-und Umweltanalytik (Eric Schweizer AG, Thun, Switzerland). A fresh batch of Changins soil was re-analyzed for pH, soil texture and soil nutrients in parallel with the field soils from Aurora and Reckenholz. Soils chemical characteristics were determined in 1:10 water (H2O, proxy for plant available nutrients) and 1:10 acetate-ammonium EDTA (AAEDTA, proxy for reserve nutrients) extracts. Total iron was analyzed by weighing 250 mg of dried and ground soil samples in 50 ml centrifuge tubes, adding 4 ml of concentrated HNO3 (65%, subboiled) and following overnight incubation at room temperature 2 ml of H2O2 (35%, trace select™) was added. Samples were vortexed for 30 seconds and then heated for extraction for 30 minutes in a microwave oven at 95°C. For analysis the samples were diluted to 50 ml and centrifuged (5’ at 1,200 rpm) and total iron (^56^Fe, ^57^Fe, analyzed in He mode) was analyzed by inductively coupled plasma mass spectrometry (ICP-MS, 7700x Agilent) using ^103^Rh and ^115^In as internal standards. Blanks were confirmed to contain <0.2% of the Fe concentrations in the samples. **Fig. S2** documents the physico-chemical characteristics of the soils at the three locations Changins, Reckenholz and Aurora.

### Sample collection

The field experiment in Reckenholz was sampled using the same protocol as in Changins (see [24] for details). In brief, shoots were removed and a 20 x 20 x 20 cm soil core containing the root system was excavated, packed in plastic bags and transported to the laboratory for sample fractionation (**Fig. S1A**). A destructive sampling was not possible in Aurora (plants were needed for another experiment) and therefore, we collected close to the base of the plants a cylinder (5 cm diameter x 20 cm depth, **Fig. S1B**) containing soil and a part of the root system. Cylinders were also packed in plastic bags and transported to the laboratory for sample fractionation.

Samples were fractionated to the different compartments in the laboratory within maximum half a day after collection on the field. The collected soil cores or cylinders were broken apart to separate the roots from the soil. The soil fraction of the cores or cylinders was thoroughly mixed and a 2 ml aliquot, representing the soil compartment, was collected and frozen at −80°C until the further analysis. The root fraction was further processed on sterile Petri plates and, using sterile scissors, we collected a 10 cm root segment corresponding to a soil depth of −5 and −15 cm. On the Petri plates, we cut the root segment into small pieces and transferred them into sterile 50 ml Falcon tubes containing 25 ml of sterile milli-Q water and we washed the rhizosphere from the roots by vigorously shaking the tubes 10 times. The roots were then transferred with sterile tweezers into fresh tubes containing 25 ml of sterile milli-Q water for an additional wash step. This was repeated 4 times and we then transferred the washed roots into 15 ml Falcon tubes to freeze-dry them for 72 hours. Lyophilized roots were then ground to a fine powder in a ball mill (Retsch GmBH, model MM301; settings 30 s at 30 Hz using on 1-cm steel ball) and aliquots, representing the root compartment, were transferred in 1.5 ml microcentrifuge tubes and stored at −80°C until further analysis. Of note, the sampling method for the root compartment does not discriminate between the inner root tissue and the root surface and therefore, we refer to the sampling unit as “root microbiota”. The rhizosphere compartment was prepared by combining all four wash fractions (4x 25 ml) using centrifugation (5 min at 3220×g, discarding the supernatant), and the resulting pellets were collected in 1.5 ml microcentrifuge tubes and stored at −80 °C until further use.

### DNA extraction, PCR and sequencing

DNA was extracted using the FastDNA SPIN kit for soil (MP Biomedical, USA) following the manufacturer’s instructions with 50-100 mg of ground roots, 100 mg rhizosphere soil, or 100 mg soil as input material. DNA concentrations were determined on a Varian Eclipse fluorescence plate reader (Agilent, USA) using the Quant-iT^TM^ PicoGreen^®^ dsDNA Assay Kit (Invitrogen, USA) and a standard solution prepared from Herring Sperm DNA (Invitrogen, USA).

Bacterial and fungal community profiles were determined following the methodology described earlier [24]. In brief, bacterial profiles are based on PCR primers 799F [38] and 1193R [39] that span the hypervariable regions V5 to V7 of the 16S rRNA gene and fungal profiles are derived from the internal transcribed spacer (ITS) region 1 amplified with PCR primers ITS1F [40] and ITS2 [41]. The **Supplementary Data 1** contains the experimental design with the sample-to-barcode assignments. PCR reactions consisted of 5-prime HotMastermix (QuantaBio, USA) (1x), bovine serum albumin (BSA, 0.3%), forward primer (200 nM), reverse primer (200 nM), with 3 ng input DNA per reaction for soil and rhizosphere samples and 9 ng for root samples, completed to 20 µl with H2O. Each sample was amplified in 3 technical replicates and one control sample without DNA per barcoded primer combination. Cycling settings were 3 minutes at 94°C for denaturation, 30 cycles (Bacteria: 30 seconds at 94°C, 30 seconds at 55°C and 30 seconds at 65°C; Fungi: 45 seconds at 94°C, 60 seconds at 50° and 90 seconds at 72°C) and a third step of 10 minutes at 65°/72°C. Reaction triplicates were pooled and confirmed for absence of contamination by running an aliquot on a 1.5% agarose gel. PCR products were then purified with a PCR clean-up kit (Macherey-Nagel, DE), quantified with the PicoGreen assay as described above, equimolarly pooled, purified and concentrated with AMpure XP beads (Beckman Coulter Inc, USA) and quantified with Qubit (Thermo Fisher, USA).

Library preparation was completed by ligation of the Illumina adapters by the Functional Genomics Center Zurich (http://www.fgcz.ch/), where they were sequenced on a MiSeq instrument in paired-end 2 × 300 bp mode (Illumina, USA). The raw sequencing data was deposited at the European Nucleotide Archive (http://www.ebi.ac.uk/ena), see **Table S1** for details linking libraries, MiSeq runs, ENA study accessions and sample IDs.

**Table 1.**
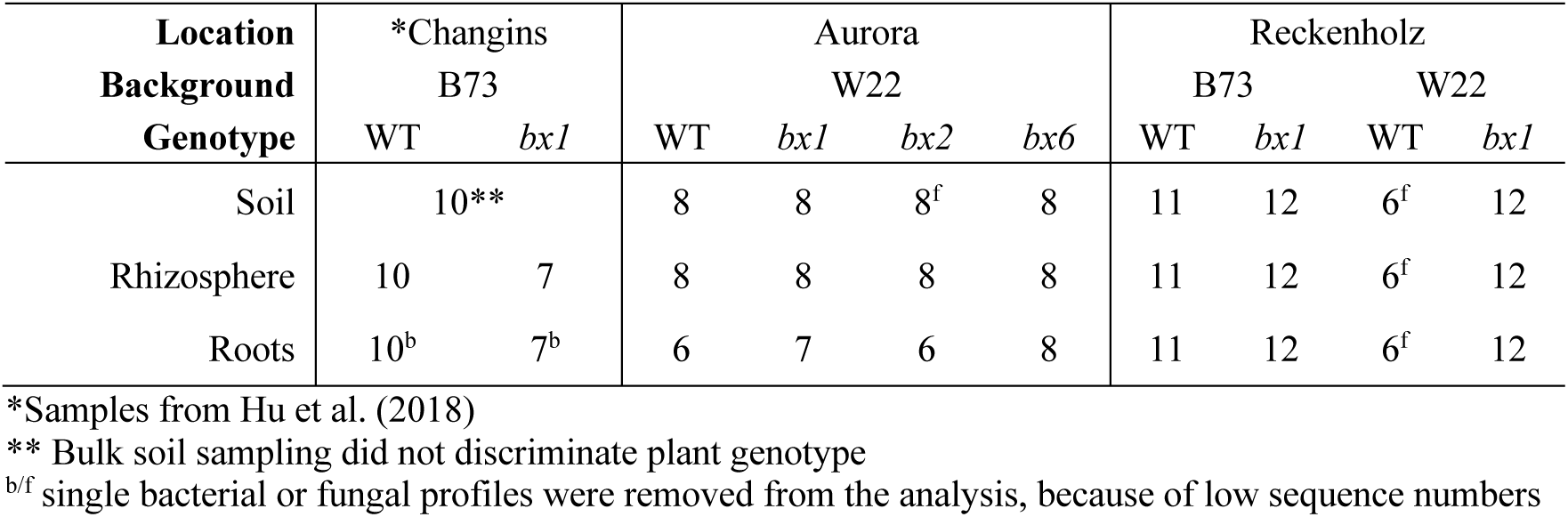
Experimental design. . Number of replicates in each sample group

### Bioinformatics

The paired-end raw reads were processed according to the bioinformatic script and parameters therein as provided in **Supplementary Data 2**. Briefly, we trimmed the low quality ends of the sequence reads R1 and R2 by cutting the reads to 280 nt and at the same time we removed all reads shorter 100 nt or reads with >1 ambiguous nucleotides using PRINSEQ (0.20.4; Schmieder & Edwards, 2011). We merged the reads with a minimum overlap of 15 nt and a maximum overlap of 250 nt with FLASH (1.2.11; Magoč & Salzberg, 2011). Individually barcoded samples were demultiplexed using Cutadapt (2.4; Martin, 2011) and filtered for GC content (range 30-70%) and quality (min. mean qual score = 20 and no ambiguous base calls) with PRINSEQ. We then used UNOISE3 (v11; Edgar, 2016a) to compute zero-radius OTUs (zOTUs) and we inferred taxonomy assignments with the Sintax algorithm [45] and the SILVA (v128, [46]) and UNITE (v7.2, [47]) databases for bacteria and fungi, respectively. We report taxonomy assignments with a >0.85 confidence cutoff. Of note, the sequence data of the field experiment in Changins [24] was reprocessed and analyzed as zOTUs in this study. **Table 1** summarizes the number of replicate microbiota profiles per compartment, field location, maize background and genotype.

### Microbiota analysis in R

All steps of the microbiota analysis were performed in R (version 3.5.1) and are documented in a markdown file that is available, together with all input files, parameter settings and functions required for replication of the analysis, as **Supplementary Data 3**. The analysis logic with the key steps are illustrated in **Fig. S3**.

The **Supplementary Data 1** contains the metadata for each sample and is loaded into R as the experimental design. Bacterial and fungal zOTUs were renamed to bOTUs and fOTUs, respectively. Microbiota profiles were filtered to exclude zOTUs classified as eukaryotes, cyanobacteria or when assigned to plant mitochondria or chloroplasts. When inspecting the mean sequencing depths of the samples, we found significant differences among our groups of samples (**Fig. S4**, Kruskal-Wallis test, *P* < 0.05). This was seen between different locations and different compartments and was especially prominent in the fungal data with up to a 6.3 fold difference in mean sequencing depth. To avoid confounding our comparisons between locations and between compartments with differences in sequencing depth, we normalized bacterial and fungal count tables by subsampling that data to 9,000 and 4,000 sequences per sample, respectively. Compared to other normalization techniques, rarefication mitigates artifacts of sampling depth more effectively, especially for datasets with low or uneven sequencing depth between groups [48]. For groups with large differences in the mean sequencing depth between groups, rarefying improves the clustering of samples according to biological origin, as well as decreasing the false discovery rate in differential abundance testing.

The rarefied count tables were utilized for alpha and beta diversity analyses using the R packages *phyloseq* [49] and *vegan* [50]. We utilized the R-package *edgeR* [51] for identification of differentially abundant zOTUs within location and plant compartment subgroups. The likelihood ratio tests were performed on data that was filtered to contain ‘quantifiable’ zOTUs (we defined ‘quantifiable’ as zOTUs with minimal abundance according to the lowest replicate number per test group (e.g., 5 replicates, min. abundance 5 sequences)) and we normalized the counts using the trimmed mean of M values method (TMM normalization, Robinson & Oshlack, 2010). Of note, TMM normalization assumes a constant abundance of a majority of species. To avoid violating this assumption and because microbes are highly variable between locations or compartments (soil, rhizosphere and roots), we first split the data by location and compartment and then applied TMM normalization within these data subsets. *P*-values were adjusted for multiple hypothesis testing [53]. For visualization, we expressed the TMM-normalized OTU abundances as percentages.

Fungal guild analysis was performed by comparing the taxonomies of the BX-sensitive fOTUs with the FUNGuild database [54]. This database uses the taxonomy of the fOTUs (done at genus or higher ranks) to assign ecological guilds (Saprotroph, Endophyte, Plant Pathogen, Animal Pathogen, Endomycorrhizal…) based on literature. We were mainly interested in the guild ‘Plant Pathogen’ and therefore, simplified the divers ecological guilds by summarizing them as ‘others’. The functional assignments are ranked with confidence categories of ‘possible’, ‘probable’ and ‘highly probable’ of which we included all.

### RESULTS

#### Sequencing effort and general overview

For this study, we determined the microbiota profiles of the two field experiments in Aurora (USA) and Reckenholz (Switzerland) by sequencing amplicons of the 16S rRNA gene and the first internal transcribed spacer region for bacterial and fungal profiling, respectively. We analyzed this new data together with existing microbiota profiles from our earlier field experiment in Changins (Switzerland; [24]). The whole analysis covers microbiota profiles of soil, rhizosphere and root compartments of WT and different mutant lines collected in Changins for the genetic background B73, in Reckenholz for B73 and W22, and in Aurora for W22 (**Table 1**). We performed the comparative microbiota analysis at the resolution of exact sequence variants, referred to as zero-radius operational taxonomic units (zOTUs), following the UNOISE method [44]. Bacterial community profiling yielded a total of 11,125,692 high-quality sequences, (range 7,541-161,133, median 41,111; **Fig. S4**). Fungal community profiling yielded a total of 3,946,277 high-quality sequences, (range 4,052-46,120, median 12,442; **Fig. S4**). Mean sequencing depths differed significantly between sampling groups and therefore, we normalized the bacterial and fungal sequence count tables by rarefying to 9,000 and 4,000 sequences per sample, respectively (see Methods). Rarefaction analysis showed that with the obtained sequencing depths, we sufficiently sampled the bacterial and fungal zOTUs (hereafter bOTUs and fOTUs, respectively) for most samples (**Fig. S5**).

To obtain an overview over the dataset, we first performed a general examination of the microbiota profiles from the different compartments and from the different locations (see **Supplementary Results**). The taxonomy, alpha- and beta diversity analyses revealed that microbiotas differed strongly between the compartments as well as between locations (**Fig. 1**, **Fig. S6-S7, Tables S2-S4**). This corroborates the work of previous studies that have shown a high context dependency of microbiotas from different compartments and/or locations [7, 16].

**Figure 1.**
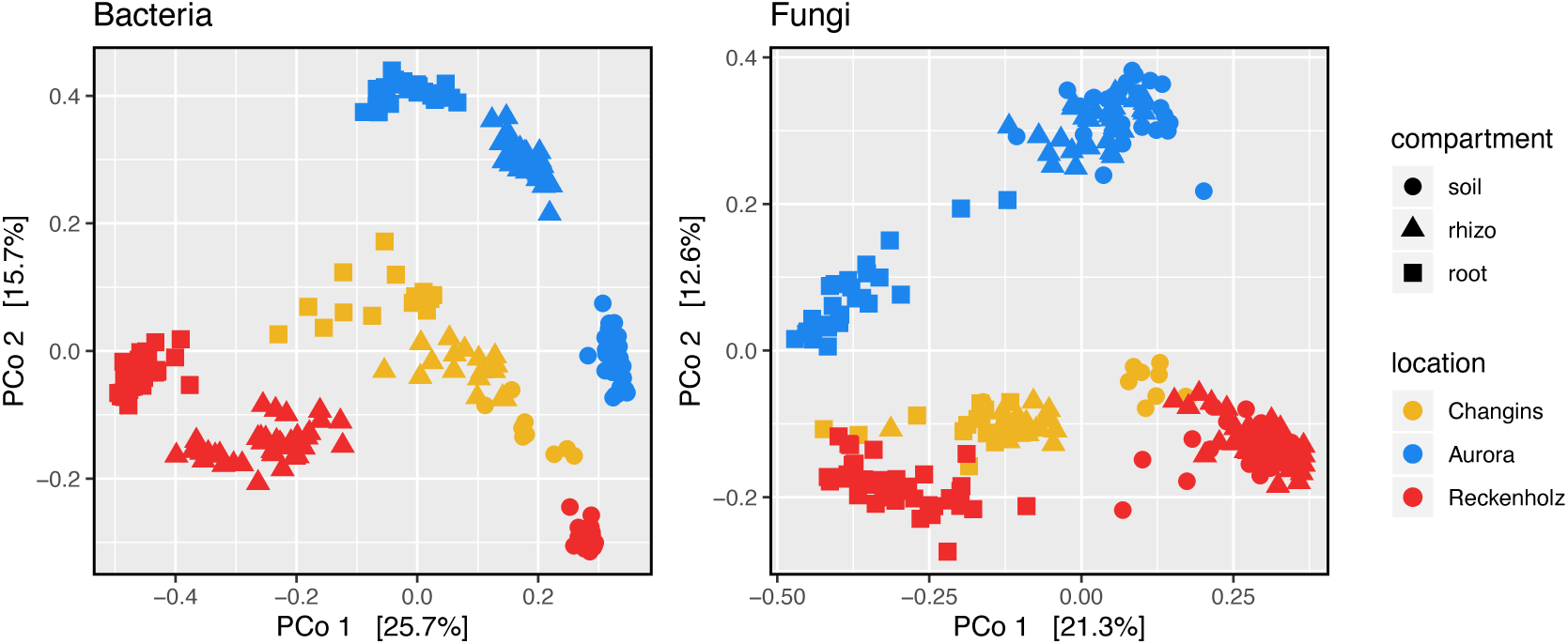
Microbiotas differ between compartments and locations. Unconstrained principle coordinate analysis (PCoA) of beta-diversity using Bray-Curtis distances of bacteria (left) and fungi (right) communities in root (circles), rhizosphere (triangles) and soil (squares) compartments from Changins (yellow), Aurora (blue) and Reckenholz (red) locations.

#### Benzoxazinoid exudation shapes rhizosphere microbial communities

To answer the first research question – How do soils, rhizosphere and roots compare in their microbiota responses to BX exudation? – we analyzed the three compartments separately. We compared the taxonomy, alpha- and beta diversities limited to WT and *bx1* mutant lines, as these plant lines were present at all locations.

In our earlier work, we had compared the rhizosphere and root communities relative to the wider soil microbiota in the field, but we did not specifically test whether BX exudation would also impact the closer soil microbiota in the soil cores that were used for the feedback experiments [24]. Therefore, we determined whether BX exudation would also impact the closer soil microbiota in the sampled soil cores of 20 x 20 x 20 cm from around WT and *bx1* plants. Comparing the phylum profiles, we found that the bacterial and fungal communities of the soil cores were similar between WT or *bx1* plants (**Table S5**). The alpha and beta diversity analyses revealed that both soil bacterial and fungal Shannon diversity were unaffected by BX exudation (**Fig. 2A**, **Tables S6 & S7**).

**Figure 2.**
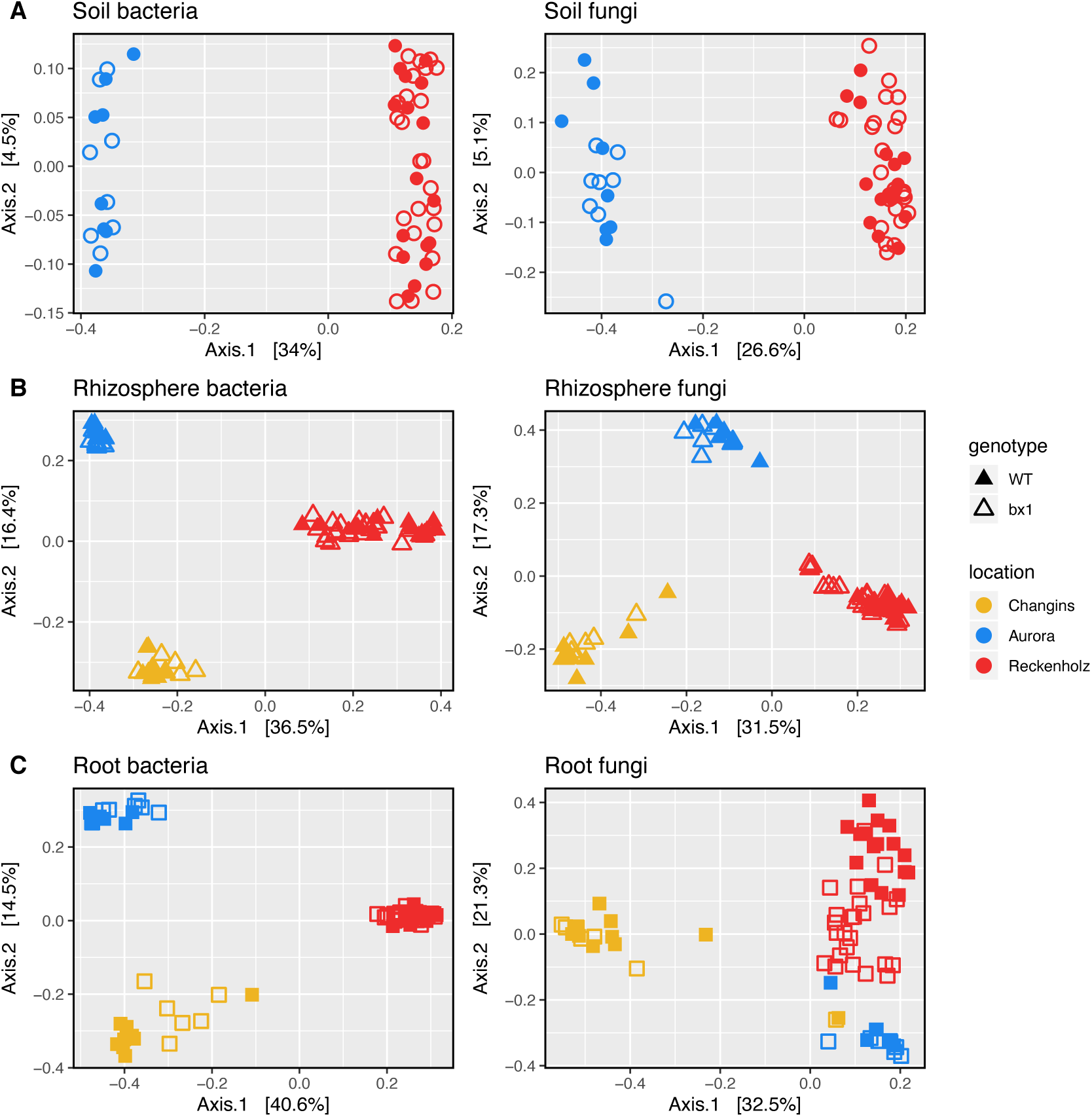
BX exudation impacts rhizosphere and root microbiotas. Compartment-wise unconstrained principle coordinate analysis (PCoA) using Bray-Curtis distances of community profiles from bacteria (left) and fungi (right). Wild-type (WT, filled) and *bx1* mutant (open) lines in soil (A, circles), rhizosphere (B, triangles) and root (C, squares) compartments from Changins (yellow), Aurora (blue) and Reckenholz (red) locations.

In the rhizosphere compartment, however, the taxonomy analysis revealed numerous bacterial and fungal phyla differing in abundance between plants that do and do not secrete benzoxazinoids (**Table S5**). In particular in Reckenholz, the rhizospheres of BX-producing plants were significantly enriched in Gammaproteobacteria (WT 56%, *bx1* 36% mean relative abundance) while the rhizospheres of BX-deficient plants contained elevated levels of Betaproteobacteria (WT 1%, *bx1* 18%), Bacteroidetes (WT 7%, *bx1* 11%), Alphaproteobacteria (WT 6%, *bx1* 11%), Deltaproteobacteria (WT 1.2%, *bx1* 1.7%) and Glomeromycota (WT 0.1%, *bx1* 0.5%), and some unassigned fungi (WT 4%, *bx1* 11%). In Aurora we only found few low abundant taxa being enriched in the rhizospheres of BX-deficient plants. Similar to the soil compartment, both bacterial and fungal alpha diversity in the rhizosphere were unaffected by BX exudation (**Table S6**). With regard to community composition, PERMANOVA quantified small yet significant effect sizes due to BX exudation of 2.2% for bacteria and 2.2% for fungi (**Fig. 2B**, **Table S7**).

In roots, the taxonomic analysis detected a few differences between WT and *bx1* plants (**Table S5**). Elevated levels of Alphaproteobacteria (WT 5.9%, *bx1* 8.8%) were found in *bx1* plants in Reckenholz. While root bacterial diversity was unaffected by BX exudation, we found an enrichment in fungal Shannon diversity in WT compared to *bx1* root samples at all three locations (**Table S6**). BX exudation significantly impacted the root microbiota, with a more pronounced effect on the fungi (6.1%) than on the bacteria (1.6%; **Fig. 2C**, **Table S7**).

In conclusion, the closer bulk soil microbiota is largely unaffected by maize BX exudation, while community composition of rhizosphere and root compartments changes significantly.

#### Effects of mutations in the BX biosynthesis pathway outweigh effects of plant genetic background

To answer the second research question – What is the influence of the plant genetic background on BX-mediated microbiota effects– we studied the WT and *bx1* mutant lines in both genetic backgrounds, B73 and W22, in Reckenholz soil. The Principle Coordinate (PCo) Analysis confirmed the major variation between *compartments* (rhizosphere vs. root, first PCo axis) and suggested that *genotype* (WT vs. *bx1*, second PCo axis) explains more variation than *background* (not apparent from the first two PCo axes), in both roots and rhizospheres (**Fig. 3A**). We then quantified the effect sizes of *background* and *genotype* relative to *compartment* employing PERMANOVA (**Table S8**). Compared to the major source of variation (*compartment* explains 35% in Bacteria and 50% in Fungi), maize *background* (2.7%; 2.5%) explained less than *genotype* (3.4%; 4.5%) in both bacterial and fungal communities. Pairwise PERMANOVA indicated for both root and rhizosphere compartments that the microbiotas differed mostly between B73 and its mutant *bx1*, whereas this differentiation was weaker in the W22 background (**Table S8**). To further characterize the impacts of *background* and *genotype* on the rhizosphere and root microbial communities, we determined the number of b/fOTUs that differed significantly between these two factors. With the exception of the rhizosphere bacteria, we found fewer b/fOTUs differing between B73 and W22, while more b/fOTUs discriminated wild-type from mutant genotypes (**Fig. 3B**, **Table S9**).

**Figure 3.**
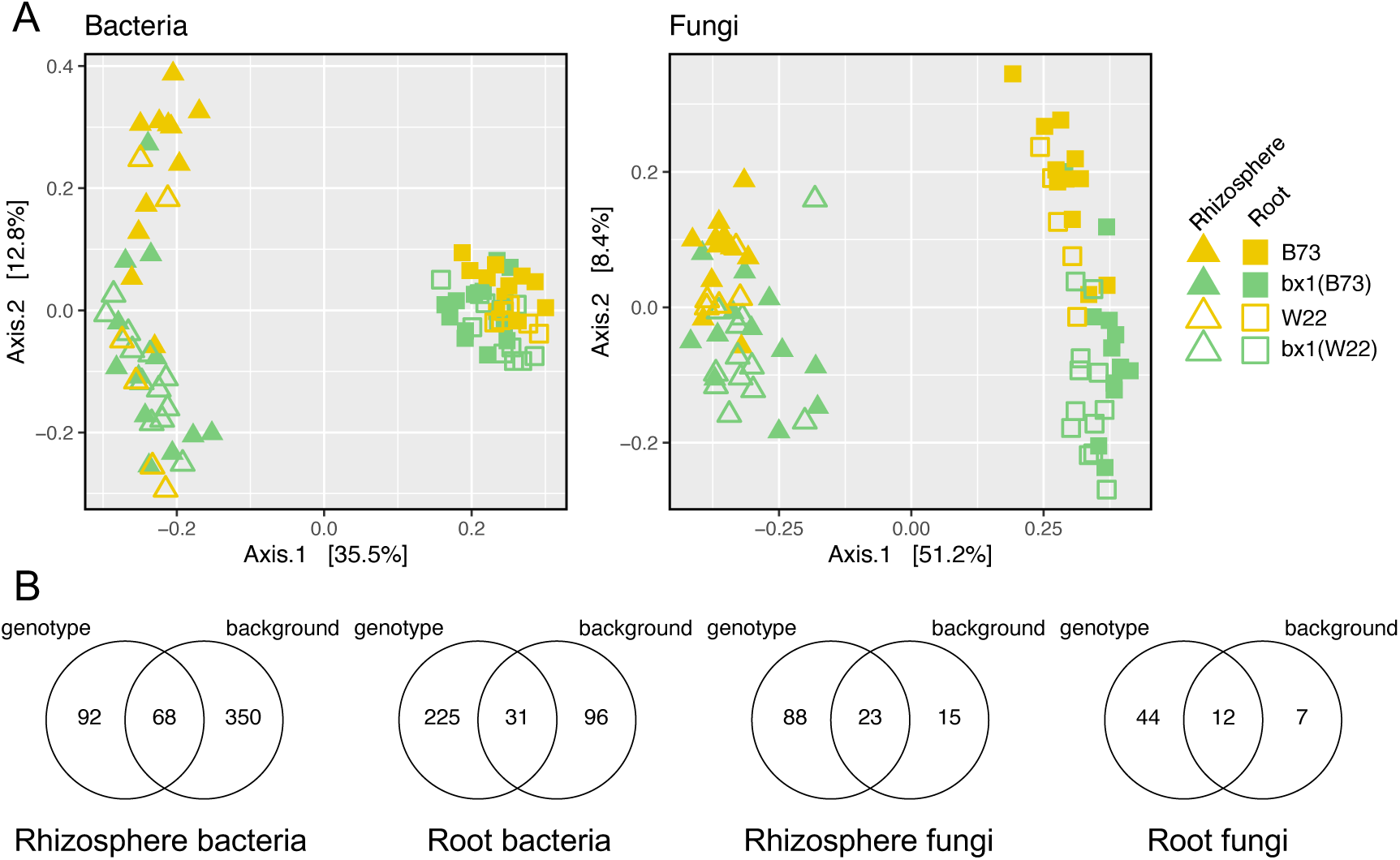
Microbiotas differ more by BX exudation than genetic background. Unconstrained principle coordinate analysis (PCoA) using Bray-Curtis distances of community profiles from bacteria (left) and fungi (right) from the Reckenholz experiment are shown in (A). Rhizosphere and root compartments discriminate as triangles and squares, respectively. Genotypes of wild-type (yellow) and *bx1* mutant (green) lines differ in color while the backgrounds B73 (filled) and W22 (open) distinguish by symbol filling. (B) Number of OTUs that differed significantly by the two factors *genotype* (WT vs. *bx1*) or *background* (B73 vs. W22) as determined by edgeR analysis (FDR < 0.05, **Table S9**).

Taken together, the effects on microbial communities comparing the two genetic backgrounds are weaker than the effects of mutations in the BX biosynthesis pathway. The effect of the mutation (WT vs. *bx*1) appears more pronounced in B73 compared to W22.

#### *BX2* and *BX6* show stronger similarity in their impact on microbiota structure compared to *BX1*

We then approached the third research question – How do different mutations in the BX biosynthesis pathway shape rhizosphere and root microbial communities? – by comparing the mutants *bx1*, *bx2* and *bx6* to the background W22, which were all grown under field conditions in Aurora. The mutant *bx2* has a BX accumulation profile similar to *bx1*, while *bx6* accumulates different BX species compared to the wild-type line W22 (see introduction).

*Compartment* explained most of the variation and separated along the first PCo axis while the second axis of the ordination indicated some clustering by *genotype* (**Fig. 4A**). Clustering by mutants was observed for rhizosphere and root bacteria, a bit weaker for root fungal communities, whereas no apparent grouping by mutants was seen among rhizosphere fungal communities. PERMANOVA quantified overall variation among the genotypes of 8.2% and 6.4% for bacterial and fungal communities, respectively (**Table S10**). Pairwise PERMANOVA revealed significant differences between wild-type and mutant plants in bacterial communities in both compartments. However, fungal communities only differed significantly between wild-type and *bx1* roots as well as between wild-type and *bx2* rhizospheres.

**Figure 4.**
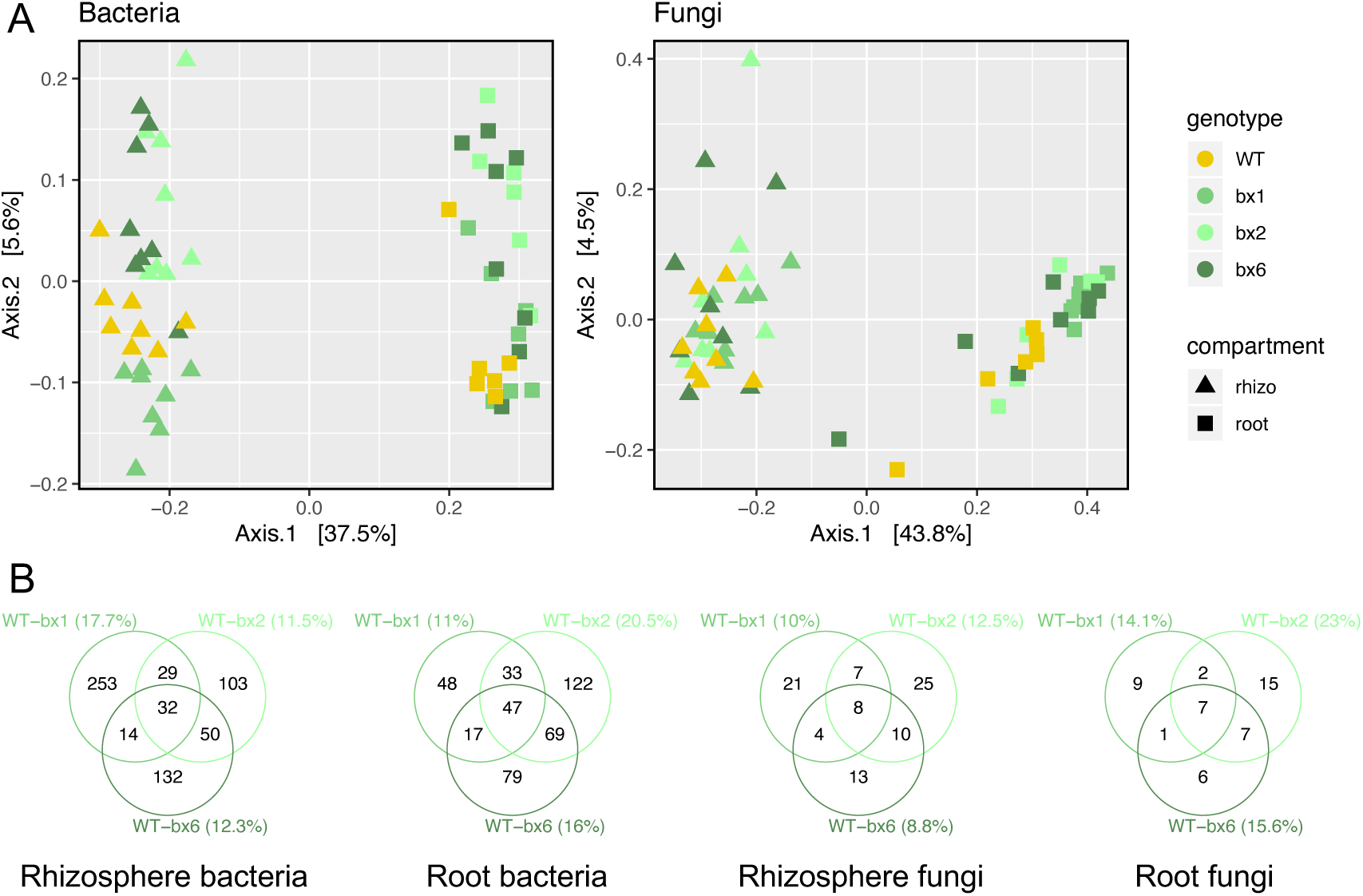
Microbiotas of *bx2* and *bx6* tend to be more similar to each other than bx1 Unconstrained principle coordinate analysis (PCoA) using Bray-Curtis distances of community profiles from bacteria (left) and fungi (right) from the Aurora experiment are shown in (A). Rhizosphere and root compartments discriminate as triangles and squares, respectively. Genotypes differ in color with wild-type W22 in yellow, the *bx1* mutant in medium, the *bx2* mutant in light and the *bx6* mutant in dark green. (B) Number of OTUs that differed significantly between the wild-type line W22 and each mutant (same colors as in A) as determined by edgeR analysis (FDR < 0.05, **Table S11**). The percentage in brackets refers to the proportion of the differentially abundant OTUs among all OTUs of the analysis to approximate an effect size of each BX biosynthesis genes for its impact on microbiota structure.

To characterize the impacts of each mutation in the BX biosynthesis pathway on the rhizosphere and root microbial communities, we determined the b/fOTUs that differed significantly between wild type and each mutant (**Table S11**). We used the proportion of differentially abundant b/fOTUs (% among all OTUs of the analysis) to approximate the impact of each BX biosynthesis gene on microbiota structure (**Fig. 4B**). Lowest effect sizes for all three mutants were seen on the rhizosphere fungi. With the exception of the rhizosphere bacteria, a lack of *BX2* appears to impact more microbes than a lack of *BX1*. And, *BX2* generally impacts more microbes compared to *BX6*. We found that the majority of b/fOTUs differed between individual mutants and the wild type, while only a small fraction of b/fOTUs discriminated all three mutants from W22 (**Fig. 4B**, **Table S11**). Inspecting the overlaps between mutants, revealed that *bx1* and *bx6* shared the lowest fraction of bacteria (rhizosphere: 14 bOTUs; root: 17), *bx1* and *bx2* an intermediate fraction (29; 33) while most of the discriminating bOTUs were found shared by *bx2* and *bx6* (50; 69). This pattern was similar with the fungi and indicates that the biosynthesis genes *BX1* and *BX6* have more distinct impacts on microbiota structure, whereas *BX2* and *BX6* have more similarities in their effects on microbiota composition than *BX1* and *BX2*.

In conclusion, the three mutants discriminated from WT plants with mostly mutant-specific sets of differentially abundant b/fOTUs. Among these b/fOTUs, we noticed the subtle pattern that *bx1* and *bx2* sharing few while *bx2* and *bx6* sharing a large part of the discriminating b/fOTUs suggesting that the BX biosynthesis genes *BX2* and *BX6* have a more similar impact on microbiota structure than *BX1*.

#### BX-sensitive microbes are location specific

To answer the fourth research question – Is there a core of microbial taxa that consistently responds to BX exudation across the different soils? – we compared the wild types and mutants grown in the 3 field locations, Changins, Aurora, and Reckenholz. We limited these comparisons to WT and *bx1* mutants, as these plant lines were present at all locations. We excluded the W22 lines at the Reckenholz location because of unbalanced sample numbers. We searched for BX-sensitive b/fOTUs –being differentially abundant zOTUs between BX producing and defective plants – using likelihood ratio tests as implemented in edgeR [51].

In roots, we found 5.6% BX-sensitive bOTUs and 9.4% BX-sensitive fOTUs at the Changins location, 11.3% bOTUs and 22.9% fOTUs in Aurora and 27.6% bOTUs and 53.7% fOTUs in Reckenholz (**Fig. 5AB**, **Table S12**). The majority of BX-sensitive bOTUs and fOTUs were specific to the each of the three locations with only a few being shared between locations. The same finding applies also to the rhizosphere compartment, where a similar location specific pattern of BX-sensitive bOTUs and fOTUs was seen (**Fig. S8, Table S12**). Comparing the BX-sensitive OTUs between rhizosphere and root samples at each location revealed that the majority were specific to each compartment while the minority was shared between them (**Fig. 5C**).

**Figure 5.**
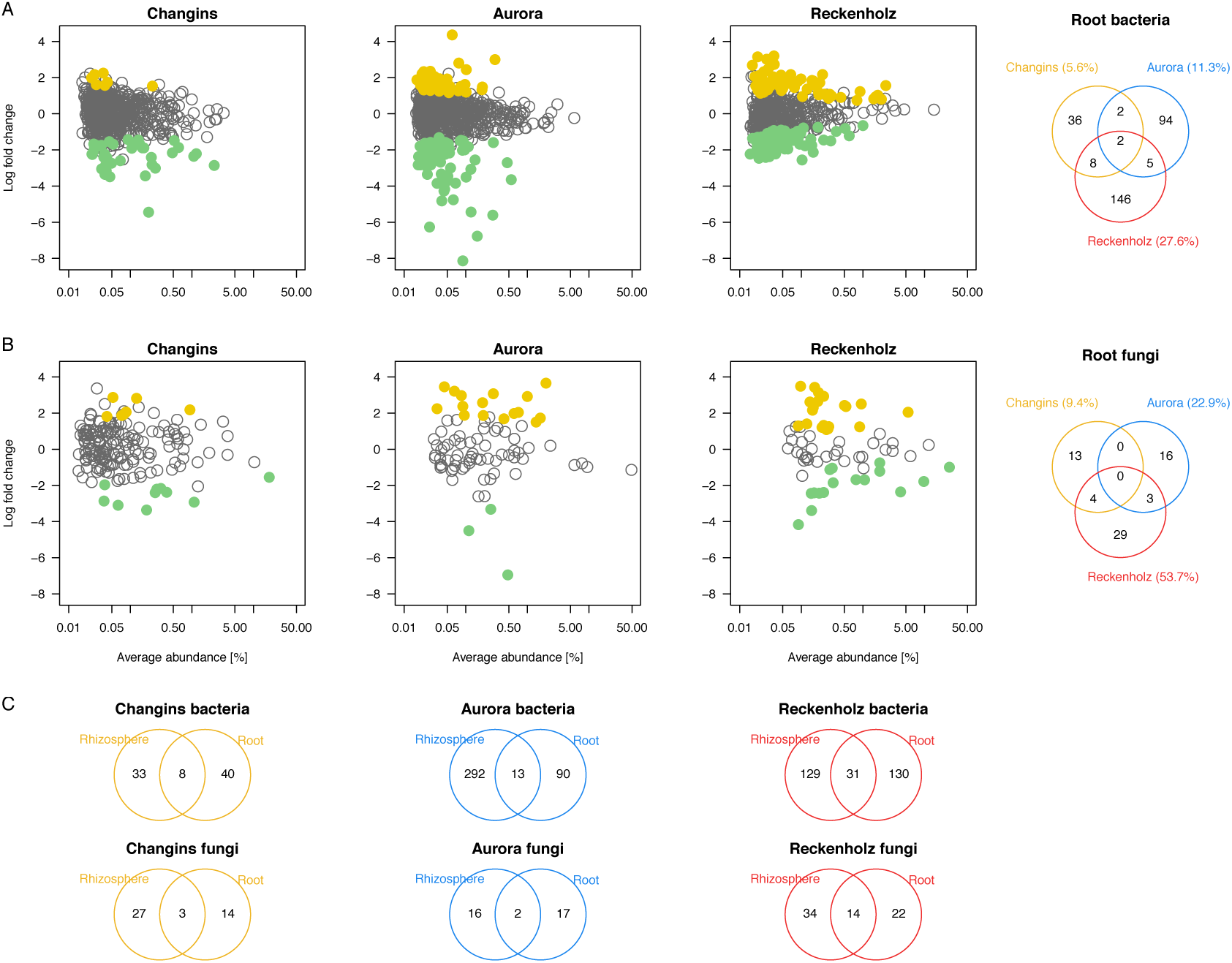
BX-sensitive root microbes are location specific The MA plots display the average abundance (in log count per million, CPM) and the log-fold change of all b/fOTUs plotted on x- and y-axes, respectively. b/fOTUs being differentially abundant between wild-type and *bx1* mutant lines (BX-sensitive OTUs) were determined by edgeR analysis (FDR < 0.05, **Table S12**). Colors refer to enriched b/fOTUs in wild-type (yellow) or *bx1* mutant (green) lines. (A) reports the root bacteria and (B) the root fungi at the locations Changins (yellow), Aurora (blue) and Reckenholz (red). The comparison of BX-sensitive b/fOTUs between locations is visualized with the Venn diagrams. (C) visualizes the comparison of BX-sensitive microbes detected in roots with the ones detected in the rhizosphere for each location (displayed in **Fig. S7**).

We then inspected these location- and compartment-specific BX-sensitive bOTUs and fOTUs for common taxonomic patterns when being enriched or depleted by BX exudation (**Fig. 6A**, **Fig. S9, Table S12**). We did not find evidence that any bacteria or fungi of certain taxonomic families that were consistently enriched by BX-producing plants in the roots or in the rhizospheres at all 3 locations. Nevertheless, we inspected, in particular, the abundance of *Methylophilaceae* in the Reckenholz and Aurora experiments, because such OTUs were previously found to be enriched by BXs in experiments conducted with Changins [24] and Sheffield soils [25]. We confirmed the BX-enrichment of a *Methylophilales* (bOTU479) and 2 *Methylophilaceae* (bOTU270 and 647) in the rhizosphere in Changins (**Fig. S10**). It is noteworthy that bOTU270 and bOTU479 were also significantly enriched in the rhizosphere samples in Reckenholz, whereas bOTU647 was significantly enriched by BXs in roots in the Aurora experiment (**Table S12**).

**Figure 6.**
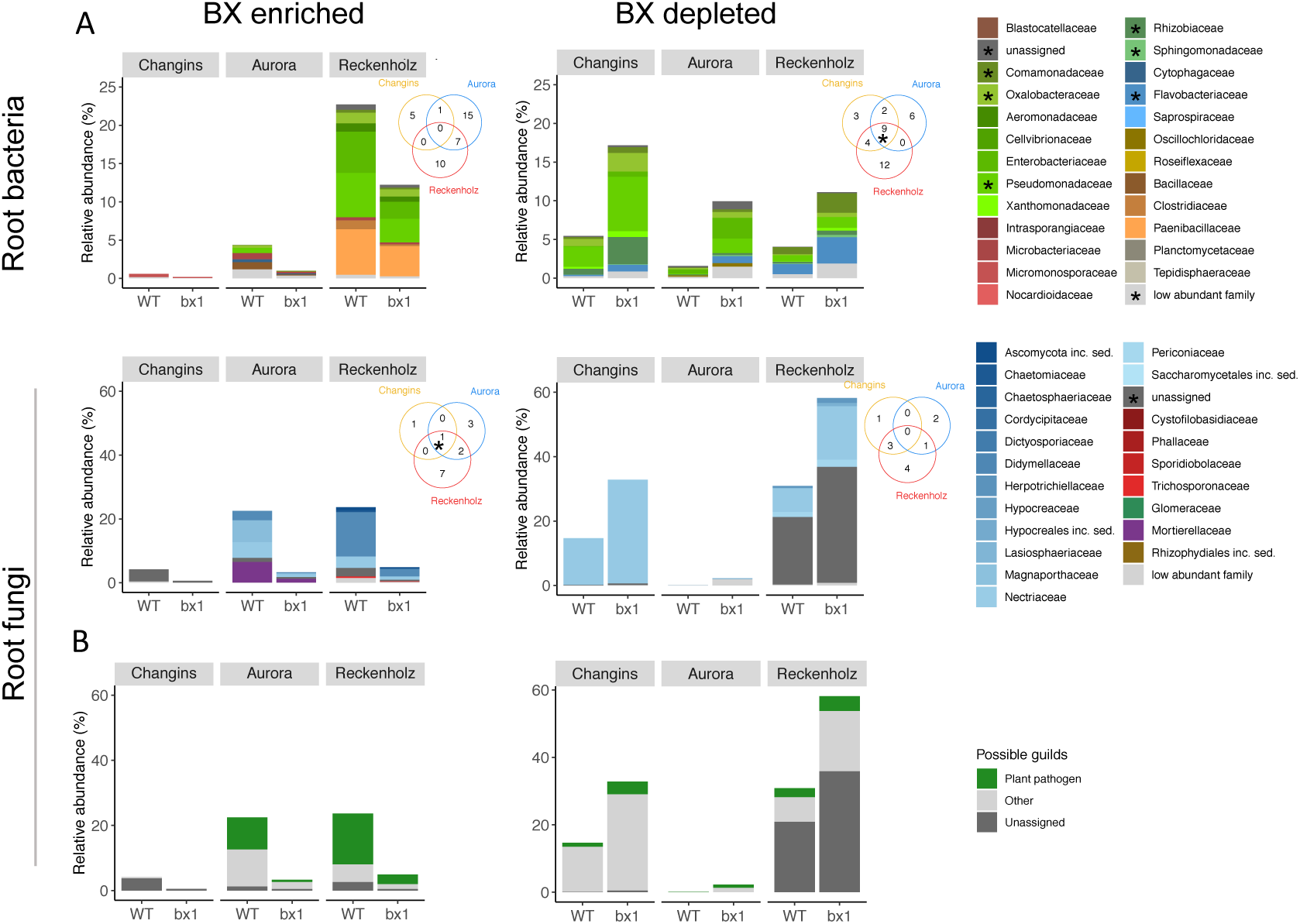
Taxonomic pattern of BX-sensitive root microbes The barplots depict for each location the mean relative abundances (in %) and taxonomies of all root bOTUs (upper panels) and root fOTUs (lower panels) that differed significantly in abundance between wild-type (WT) and *bx1* mutant lines (i.e., the BX-sensitive b/fOTUs as determined by edgeR analysis, FDR < 0.05, **Table S12**). The BX-enriched (left panels) and BX-depleted taxa (right panels) correspond to the same yellow (enriched in WT) and green (enriched in *bx1*) b/fOTUs of Fig. 5A, respectively. Individual b/fOTUs are displayed in a stacked manner sorted by their taxonomic assignment at family level. The Venn diagram insets compare the family assignments of the BX-sensitive taxa between the locations Changins (yellow), Aurora (blue) and Reckenholz (red). Overlapping family assignments are indicated with asterisks in the plot and marked in the taxonomy legend. (B) visualizes the proportion of assignments to ‘plant pathogen’ among the FUNGuild annotations. The sets of BX-enriched and BX-depleted root fOTUs from each location were annotated individually to their ecological guilds.

While a taxonomic pattern was not found among BX-depleted fungi, we found that the bOTUs of same taxonomic families were consistently depleted at all 3 locations from roots of BX producing plants (**Fig. 6A**). BX-depleted root bacteria were commonly assigned to families including the *Pseudomonadaceae*, *Oxalobacteraceae*, *Flavobacteriaceae*, *Rhizobiaceae*, *Comamonadaceae* and *Sphingomonadaceae*. In agreement with this observation, *Flavobacteriaceae* and *Comamonadaceae* were also consistently depleted in rhizosphere samples of WT plants at all 3 locations (**Fig. S9A**). Aside of this common taxonomic pattern of BX-depleted bacteria, most of the BX-dependent enrichments and depletions were location-specific (**Table S12**, see **Supplementary Results**).

Because of earlier reports [25, 26], we investigated whether microbiotas of BX producing plants would comprise fewer fungal species with taxonomic links to plant pathogens. We used FUNGuild to annotate the fOTU data to ecological guilds [54], and identified fOTUs with possible assignments to plant pathogens among the fOTUs that were BX-enriched or BX-depleted. Over all three locations, while we found only few and non-abundant BX-sensitive fOTUs in the rhizosphere samples (**Fig. S9B**), we observed in roots more fOTUs with possible links to plant pathogens and they were particularly abundant on BX producing plants in Reckenholz and Aurora (**Fig. 6B**). This observation suggests that wild-type maize plants attract fungal pathogens by BX exudation.

Taken together, the exudation of BXs affects different sets of bacteria and fungi in the different soils and in the different compartments. While BX producing plants did not consistently enrich microbes of certain taxonomic lineages in their roots or rhizospheres, we found that BX exudation depleted bacteria assigned to *Flavobacteriaceae* and *Comamonadaceae* from root and rhizosphere compartments as well as appeared to enrich for plant-pathogenic fungi.

##### DISCUSSION

In our earlier work, we showed that the secretion of BXs by a first generation of maize plants can drive microbial feedbacks on growth and defense in the next generation of maize plants [24]. The mechanistic basis for these feedbacks was that the secretion of BXs altered the structure of root and rhizosphere microbial communities. Such BX impacts on microbiotas were confirmed in young plants [25] and in semi-artificial rhizobox systems [26]. To study conserved microbiota response patterns to BX exudation we profiled soil, rhizosphere and root microbial communities in two new field experiments in Aurora and Reckenholz that we conducted similar to the first one in Changins [24].

#### Genetic background

Microbiota profiles of the WT and *bx1* mutant plants differed in the backgrounds B73 [24] and W22 [25, 26]. We grew B73 and *bx1*(B73) in parallel with W22 and *bx1*(W22) to learn ‘How the genetic background and the mutation in BX1 compare in their effects on microbial communities?’. While B73 and W22 differ at a multitude of genetic loci, the genetic variation between wild-type and mutant lines is low, as they differ mainly in the *BX1* gene, which defines their in-/ability of BX synthesis. The *bx1* mutant in B73 represents a near-isogenic line based on five backcrosses in this genetic background [35], while *bx1*(W22) contains a single *Ds* transposon insertion compared to W22 [37]. Ordination analysis (**Fig. 3A**), variation partitioning (**Table S8**) as well as the number of differentially abundant OTUs between these two factors (**Fig. 3B**, **Table S8**) revealed that the presence/absence of a functional *BX1* gene had more impact on roots and rhizosphere microbiotas than multi loci genetic differences between the backgrounds B73 and W22. Multiple genetic differences, such as between lines of different backgrounds, plant varieties, or accessions, typically account for ∼5-6% of variation in root and rhizosphere microbiota composition [7, 16]. Since, we found a similar level of microbiota variation (**Table S8**) comparing plants that are genetically nearly identical but differ in a discrete chemical pathway, this suggests that the BXs exudation presents a key trait by which maize plants structure their root and rhizosphere microbiota.

The effect of the mutation (WT vs. *bx1*) appeared more pronounced in B73 compared to the background W22 (**Table S8**). The multiple genetic differences between the two backgrounds serve as the obvious explanation for this observation whereas, mechanistically, this could be due to quantitative differences in the amounts of BXs that are exuded by these two backgrounds. The W22 background produces about 15% lower levels of total BXs in the roots compared to B73 [31] and therefore, possibly structures its root and rhizosphere microbiota to a lesser extent than B73 does. Specific work is needed to validate quantitative BX effects on microbiota composition, for instance, by studying BX overexpression lines [55].

#### Microbiotas of BX biosynthesis mutants

Studying the broader impact of BXs exudation on the root and rhizosphere microbiota, we hypothesized that variations in microbiota composition will be consistent with the BX accumulation patterns of the mutants. The mutants *bx1* and *bx2* secrete very low levels of BXs whereas *bx6* secretes different BXs (see introduction for details). Consistent with our hypothesis, the microbiota profiles of BX-producing WT plants would differ most strongly compared to the mutants *bx1* and *bx2,* and the profiles of the two mutants would share similarities with each other. We then saw two scenarios for *bx6* in this hypothesis: a) if the total amount of secreted BXs matters, the microbiota profiles of *bx6* would be like WT plants or b) if the speciation of BXs is relevant, *bx6* would display an intermediate or different microbiota composition compared to WT, *bx1* and *bx2* plants.

We found mutant-characteristic profiles for rhizosphere and root bacteria, a weak genotype signature for root fungi and no effect on rhizosphere fungal communities (**Fig. 4A**, **Table S10**). The analysis of the number of differentially abundant b/fOTUs (each mutant vs. WT) and their overlap between mutants revealed that the biosynthesis genes *BX1* and *BX6* have the least and *BX2* and *BX6* share most similarities in their effects on microbiota composition (**Fig. 4B**, **Table S11**). Hence, we rejected the first part of the aforementioned hypothesis (Most overlap expected between *BX1* and *BX2*) and concluded that factors additional to the BX accumulation pattern explain the microbiota profiles between the mutants. Regarding the second part of the hypothesis, we concluded that because the microbiota of *bx6* was found to be different compared the wild-type plants, that the speciation of BXs is relevant for root and rhizosphere microbiota assembly.

Coming back to additional factors than BX accumulation which could explain the microbiota profiles between the mutants: Firstly, BX precursors could accumulate differently in roots of the *bx1*, *bx2* and *bx6* mutant plants. Of these, volatile indole, which is produced by the indole-3-glycerolphosphate lyase IGL independently of BX1 is known as a potent signaling molecule in cereals [56–58]. Volatile indole has so far only been found in maize leaves; it remains undetectable in maize roots [58], despite transcriptional activity of IGL [59], possibly because it is rapidly metabolized at the root surface. While *bx1* mutants may show enhanced production of IGL-dependent volatile indole through increased precursor availability, *bx2* mutants may show enhanced production of BX1-dependent indole due to a lack of oxidation to oxindole. The prevalence and impact of indole in maize roots remains to be uncovered.

Secondly, benzoxazinoids may regulate the production of other secondary metabolites. Wheat plants that overexpress a maize methyl-transferase which shifts benzoxazinoid production from DIMBOA-Glc to HDMBOA-Glc also accumulate higher amounts of ferulic acid in the leaves [60]. Furthermore, Cotton *et al*. (2019) found differences in metabolic features in the roots of *bx1*, *bx2* and *bx6* mutants that could not be linked to BXs, but matched m/z values of flavonoids. Correlation analysis between microbes and metabolites revealed that, although the abundance of BX-stimulated OTUs correlated positively with BXs, the same OTUs also correlated positively with other features. Half of the features which correlated positively with BX-stimulated OTUs were assigned to potential flavonoids. Given that flavonoids can function as a semiochemical for root bacteria [61], it was suggested that these metabolites, possibly jointly with BXs, structure the root microbiota. More work is required to assess the contribution of BX-dependent metabolites to structuring root microbiota.

With respect to the mutants *bx1*, *bx2* and *bx6* in the W22 background, there is the particularity that the genes at the transposon launch sites participate in the flavonoid pathway. Both launch sites affect the downstream anthocyanin synthesis, but while R1 (*bx1* and *bx6* lines) is a bHLH transcription factor that could be acting pleiotropically, A1 (launch site for *bx2* line) is a structural gene downstream in the pathway [62]. However, the genetics of these launch sites is unlikely to drive BX-dependent microbiota effects, as a differential assembly was also found in the B73 background, which is unaffected in the flavonoid pathway. Our results are congruent with the hypothesis that BXs may act directly as well as indirectly via precursors or other metabolites to shape rhizosphere microbiota.

#### BX-sensitive microbes in different soil types

We expected the rhizosphere and root microbiotas to be different between the three tested field locations because soil-type - referring to the specific biogeochemical properties of a soil - is a well-known primary determinant of microbial community structure [7, 16]. We were interested to test whether, beyond soil-type-dependent microbiota specificities, there is a core of microbial taxa that responds to BX exudation across the different soils. However, we found most BX-sensitive microbes, whether enriched or depleted, to be location-specific (**Fig. 5**, **Table S12**). This was already seen earlier when comparing the BX-sensitive taxa in Changins soil [24] to the ones detected in Sheffield soil [25]. Considering the location-specificity of BX-effects on the root and rhizosphere microbiota, it appears plausible that the exudation quantities of BXs also may vary depending on the local biogeochemical environment at different field locations. We hypothesize that an environmentally regulated exudation of BXs could be linked to the phytosiderophore function of BXs, as these root-secreted compounds complex iron for plant uptake [31]. We need further work to clarify whether maize plants adjust the levels of BX exudation in a context-dependent manner and eventually in response to the levels of available iron in soil.

The finding that BX-sensitive microbes are location-specific, with only a few being shared between soils (**Fig. 5**, **Table S12**), rules out that BXs would selectively enrich or deplete the same microbial species across different soil types. Although they are not enriched consistently at each location and in each compartment, we found a significant enrichment of *Methylophilaceae* in rhizosphere samples in Reckenholz and in roots in Aurora from BX producing plants (**Fig. S10**, **Table S12**), similar to the experiments conducted with soils in Changins [24] and Sheffield [25]. The detection of the same sequence variants (at the level of individual zOTUs) in experiments with different soils on European and American continents makes it unlikely that it derived from soil and points to the possibility that these bacteria originate from the maize seed material. Some methylotrophic bacteria such as *Methylobacteria* (*Methylobacteriaceae*, being Alphaproteobacteria) or *Methylophilus* (*Methylophilaceae*, being Betaproteobacteria) have been reported as maize seed endophytes [63–66]. Future experiments, e.g. by sequencing kernels or roots from plants grown in sterile substrate or culturing seed endophytes on media with methanol as sole carbon source, are needed to clarify the endophytic presence of *Methylophilaceae* bacteria in seeds of BX producing maize lines.

When inspecting the BX-sensitive microbes at higher taxonomic rank, we noticed that BX-producing plants consistently depleted certain microbe families from their roots or rhizospheres. BX-producing plants mainly depleted bacteria assigned to the *Flavobacteriaceae* and *Comamonadaceae* from their roots or rhizospheres (**Fig. 6**). A similar pattern was seen among the few BX-sensitive OTUs in 17-day-young plants, with the majority being depleted while a smaller fraction was enriched by BXs [25]. Of note, only the fraction that was depleted included *Flavobacteriaceae* and *Comamonadaceae*. Besides depletion of *Flavobacteriaceae*, they also found that other Bacteroidetes bacteria such as *Cytophagaceae* and *Chitinophagaceae* were negatively affected by BX exudation. We also noticed in our dataset that *Cytophagaceae* were depleted by BX exudation in Changins and Aurora locations (**Table S12**). Together with the observation that bacterial OTUs assigned to the phylum Bacteroidetes tended to be depleted by BX exudation also in an earlier study [26], suggests an overall negative impact of BX exudation on Bacteroidetes bacteria.

While we did not observe that BX-producing plants would consistently enrich certain taxonomic groups of microbes to their roots or rhizospheres, we found evidence for an enrichment of potential fungal pathogens in the FUNGuild analysis (**Fig. 6B**). We observed more fOTUs in roots with possible links to plant pathogens and they were particularly abundant on BX-producing plants in Reckenholz and Aurora. As this is in contrast of earlier findings [25, 26], our observation that wild-type maize plants may attract fungal pathogens by BX exudation requires further examination also with other approaches such as e.g. shotgun metagenomics.

The enrichment of potential fungal pathogens and the depletion of *Flavobacteria* and other Bacteroidetes bacteria by BX exudation is interesting considering that sugar beet plants specifically enrich bacteria of two genera of the Bacteroidetes in response to fungal infection [67]. Carrion *et al.* (2019) reported that *Chitinophaga* and *Flavobacteria* have characteristic chitinase genes and previously unknown biosynthetic gene clusters encoding secondary metabolites that are essential for disease suppression. Consistent with their discovery we also observed an antagonistic relationship between Bacteroidetes bacteria and fungal pathogens: When we found an enrichment of Bacteroidetes bacteria there were low amounts of potentially pathogenic fungi (**Fig. 6**, roots and rhizospheres of *bx1* plants). In contrast, when Bacteroidetes bacteria were low in abundance we noticed an enrichment of potentially pathogenic fungi (roots and rhizospheres of WT plants). Future work is needed to disentangle whether direct microbe-microbe interactions govern this antagonistic relationship or whether the BXs function as driving force with possible repellent and attractive activities on Bacteroidetes bacteria and pathogenic fungi, respectively.

#### BX effects on microbiota in soil cores

In addition to the research questions on genetic background, mutants, and soil type, we wanted to close the knowledge gap whether the microbiota in the soil cores might also be affected by BX exudation. This question remained open from our earlier work, where we had compared the rhizosphere and root communities relative to the soil microbiota at field scale, which was analyzed based on random soil samples collected from across the field [24]. However, the microbial feedback experiments were conducted using 20 x 20 x 20 cm soil cores, representing a closer soil microbiota, and we had not specifically tested this soil core compartment for eventual effects of BX exudation. Therefore, we analyzed DNA samples from such soil cores taken in the fields of Reckenholz and Aurora and profiled this closer soil microbiota. We found that this closer soil microbiota was largely indistinguishable, irrespective of whether the soil cores were collected from WT or *bx1* plants (**Fig. 2A**, **Tables S5-S7**). This finding suggests that the amount of BXs, which was secreted into a 20 x 20 x 20 cm soil volume, becomes diluted to such an extent that BX-dependent microbiota differentiation is no longer seen. Nevertheless, we know that the overall population of microbes in the soil cores provokes the robust feedbacks on a next generation of maize [24]. Our interpretation is that, although no BX-dependent microbiota differentiation is seen at the scale of whole soil cores, such cores probably contain aggregates of former BX rhizospheres, as well as also root fragments from the previous plant generation, and that these BX microbiota hotspots suffice to trigger the observed feedbacks.

#### CONCLUSIONS

Our findings that exuded BXs function in controlling the abundance of Bacteroidetes bacteria and suppressing fungal pathogens together with the positive feedback effects on plant health [24], positions the BX pathway as an interesting target to maintain or further enhance in breeding programs. Incorporating plant-microbiome interactions into breeding programs relies on plant loci that explain heritable traits of plant microbiomes. The big challenge remains to identify plant genes that - beyond the strong environmental influences by soil type, climate, or field management - are responsible for the abundance of certain beneficial taxa or the expression of beneficial microbiome traits. This is a demanding task, as the heritability of plant microbiota composition is notoriously low [7, 16]. Even the most comprehensive analysis of plant microbiota heritability, although identifying a few taxa with genetically explained abundance, concluded that their heritabilities were low [68, 69]. This analysis included close to 5000 maize rhizosphere microbiota profiles from the 27 genetically diverse maize inbred lines, which were planted in partly repeated field experiments at three sites to include variation in environment and time. Comparisons between genetic lines, plant varieties or accessions, which differ from each other genetically by a multitude of allelic variants, typically account for ∼5-6% of variation in microbiota composition [7, 16]. We found a similar level of microbiota variation comparing plants that are genetically nearly identical but differ in a discrete major chemical pathway (**Table S8**). We argue that studying candidate root exudate pathways was a more promising approach to find host loci of heritable plant microbiota members compared to unbiased genetic diversity screens.

#### DECLARATIONS

##### Ethics approval and consent to participate

Not applicable

#### Consent for publication

Not applicable

#### Availability of data and material

The raw sequence data generated of this study are available from the European Nucleotide Archive repository [http://www.ebi.ac.uk/ena, see **Table S1** for ENA study accessions and sample IDs]. All analysis code to reproduce this study is included in this published article [**Supplementary Data 2** contains the bioinformatic script and **Supplementary Data 3** documents the microbiota analysis in R].

#### Competing interests

The authors declare that they have no competing interests

#### Funding

This study was supported by the Swiss State Secretariat for Education, Research and Innovation (grant C15.0111 to ME, MvdH and KS), the University of Bern through the Interfaculty Research Cooperation ‘One Health’, and US National Science Foundation award 1339237 to GJ.

### Authors’ contributions

KS, ME and MvdH conceived the original project and KS supervised the experiments. SC and KS designed the experiments. GJ and MvdH provided field experiment infrastructure. GJ and ME provided seeds. SC performed the experiments and the molecular work. HG and MB performed soil analyses. JCW and SC performed the bioinformatic analysis. SC and KS analyzed the data. SC and KS wrote the manuscript with input from all authors.

## Acknowledgements

We thank Juerg Hiltbrunner (Agroscope), Fritz Kaeser (Agroscope) and Kevin Ahern (Boyce Thompson Institute) for support with the field experiments. We thank Dr. Lucy Poveda from the Functional Genomics Center in Zürich for technical support in MiSeq-sequencing.

## SUPPLEMENTARY INFORMATION

### SUPPLEMENTARY RESULTS

#### Analysis of ‘compartment’ and ‘location’ effects

We examined the microbiota profiles for ‘compartment’ and ‘location’ effects by analyzing the dataset based on taxonomy, alpha- and beta diversity.

#### Community taxonomy differs by locations and compartments

In a first step, we examined the taxonomy profiles at phylum level (**Table S2**). On average over all locations, the bacterial root microbiota was significantly enriched in Gammaproteobacteria (soil 7%, rhizosphere 26%, roots 33%), Actinobacteria (soil 10%, rhizosphere 6%, root 17%), Alphaproteobacteria (soil 8%, rhizosphere 14%, roots 13%) and Firmicutes (soil 0.5%, rhizosphere 1%, root 5%, **Fig. S6**). In contrast, Chloroflexi (soil 10%, rhizosphere 3%, root 2%), Deltaproteobacteria (soil 7%, rhizosphere 3%, root 1%), Acidobacteria (soil 7%, rhizosphere 2%, root 0.3%), Verrucomicrobia (soil 6%, rhizosphere 2%, root 0.2%) and Gemmatimonadetes (soil 7%, rhizosphere 1%, root 0.3%) were significantly more abundant in soil (**Fig. S6**). While the taxonomic composition differed little between soils from different locations, we noticed more variation in the rhizosphere and root compartments. The rhizosphere samples in Reckenholz had a high proportion of Gammaproteobacteria, while Changins and Aurora were characterized by pronounced Bacteroides and Betaproteobacteria, respectively. In roots, high proportions of Actinobacteria were found in Changins, whereas Gammaproteobacteria were abundant in Reckenholz and Alphaproteobacteria were typical for Aurora (**Fig. S6**).

For the fungal microbiota, over all locations Ascomycetes (soil: 52%, rhizosphere: 51%, root: 86%) and Basidiomycetes (soil: 6%, rhizosphere: 8%, root: 8%) were significantly more abundant in roots (**Fig. S6**, **Table S2**). Soil fungal communities were significantly enriched in Mortierellomycota (soil: 20%, rhizosphere: 21%, root: 2%) and Glomeromycota (soil: 5%, rhizosphere: 1%, root: 2%) and they differed only marginally between locations. Rhizosphere communities in Aurora and Reckenholz also contained abundant Mortierellomycota whereas Basidiomycota were the second most abundant fungi in Changins. Root fungal communities were dominated by Ascomycetes at all locations (Changins: 61%, Aurora: 64%, Reckenholz: 64%). Some Basidiomycota and Glomeromycota fungi were also found enriched in roots from Changins, while they were not much detected in Reckenholz and Aurora (**Fig. S6**).

Taken together, the taxonomic composition of the microbiotas differs strongly between soil, rhizosphere and roots as well as between the locations Changins, Aurora and Reckenholz.

#### Community diversity varies by compartment and is location specific

To investigate alpha diversity of the microbial communities, we rarefied the bacterial and fungal datasets to even sampling depths of 9,000 and 4,000 sequences, respectively, and statistically compared the Shannon metrics between compartments and locations (**Table S3**). Consistent with the rarefaction analysis (**Fig. S5**), bacterial diversity decreased from soil to rhizosphere to roots in all locations, was similar in the soil compartment at all three locations, but was found to be systematically lower in the rhizosphere and root compartments in Reckenholz soil (**Fig. S7**). Similarly, fungal diversity was high in soil and rhizosphere compartments and decreased in roots at all three locations (**Fig. S7**). There is notably high rhizosphere diversity specific to the Aurora location. We concluded that the detected alpha diversity varied strongly between compartments and displayed location specific patterns.

#### Community composition differs by location and compartment

Finally, we assessed microbial β-diversity by factors ‘compartment’ and ‘location’ using permutational multivariate analysis of variance PERMANOVA and unconstrained principle coordinate (PCo) analysis. The R^2^ values of PERMANOVA were taken to approximate effect sizes of the different factors. They indicated that overall ‘compartment’ explained most of microbial community variation (Bacteria 25.6%, Fungi 19.7.5%), followed by ‘location’ (23.2%, 21.8%; **Table S4**). Unconstrained principle coordinate (PCo) analysis visualizes this conclusion with the samples largely separating these two factors along PCo axes 1 and 2, respectively (**Fig. 1**). For fungi, the pattern is similar except that the two PCo axes explained slightly less of the overall variation. This confirms that the composition of microbial communities strongly differ between compartments and different locations.

### Analysis of BX-sensitive microbes

We have reported the common taxonomic patterns of BX-depleted microbes in the main text and here we detail the location-specific enrichments and depletions of BX-sensitive microbes (**Table S12)**.

Globally, there were more BX-depleted than BX-enriched b/fOTUs, more of the BX-sensitive b/fOTUs were seen in the rhizosphere than in roots, and more bOTUs were BX-sensitive than fOTUs. On roots, diverse low abundant bacteria were enriched on BX producing plants at all 3 locations while a specific enrichment of abundant *Paenibacillaceae* (2 bOTUs), a *Pseudomonadaceae* and an *Enterobacteraceae* was seen only in Reckenholz. Abundant BX-enriched fOTUs were *Nectriaceae*, *Mortierellaceae, Gaeumannomyces* in Aurora and *Didymellaceae* (Reckenholz) whereas several low abundant fOTUs were found in Changins. In contrast, BX exudation depleted the following bOTUs from WT roots: abundant *Pseudomonadaceae* and *Rhizobiaceae* were depleted in Changins and several low bacteria were depleted in Aurora and Reckenholz. BX-depleted root fungi included abundant *Nectriaceae* (Changins and Reckenholz), *Herpotrichiellaceae* and several *Pleosporales* (Reckenholz) while in Aurora diverse low abundant fungi were depleted.

In the rhizospheres of BX-producing plants, mostly low abundant bacteria and in particular *Pseudomonas* (Reckenholz) were enriched (**Table S12)**. We noticed that in Reckenholz, BX exudation enriched for diverse fOTUs, the abundant ones belonging to *Mortierellaceae*, *Chaetomiaceae* and *Saccharomycetales* besides an unassigned fOTU. In Changins specifically, enriched fOTUs included several *Glomeraceae* along with an *Agaricales*. Conversely, low abundant bacteria were depleted in rhizospheres including *Flavobacteriaceae* (Changins and Aurora), *Saprospiraceae* (Aurora and Reckenholz) and several location-specific groups (*Burkholderiales* in Changins, *Blastocatellaceae* in Aurora and *Sphingomonadaceae* and *Xanthomonadaceae* in Reckenholz). Low abundant (Aurora) or several abundant unassigned or diverse fOTUs were depleted by BX-exudation in Changins and Reckenholz, respectively.

## SUPPLEMENTARY FIGURES

**Figure S1.**
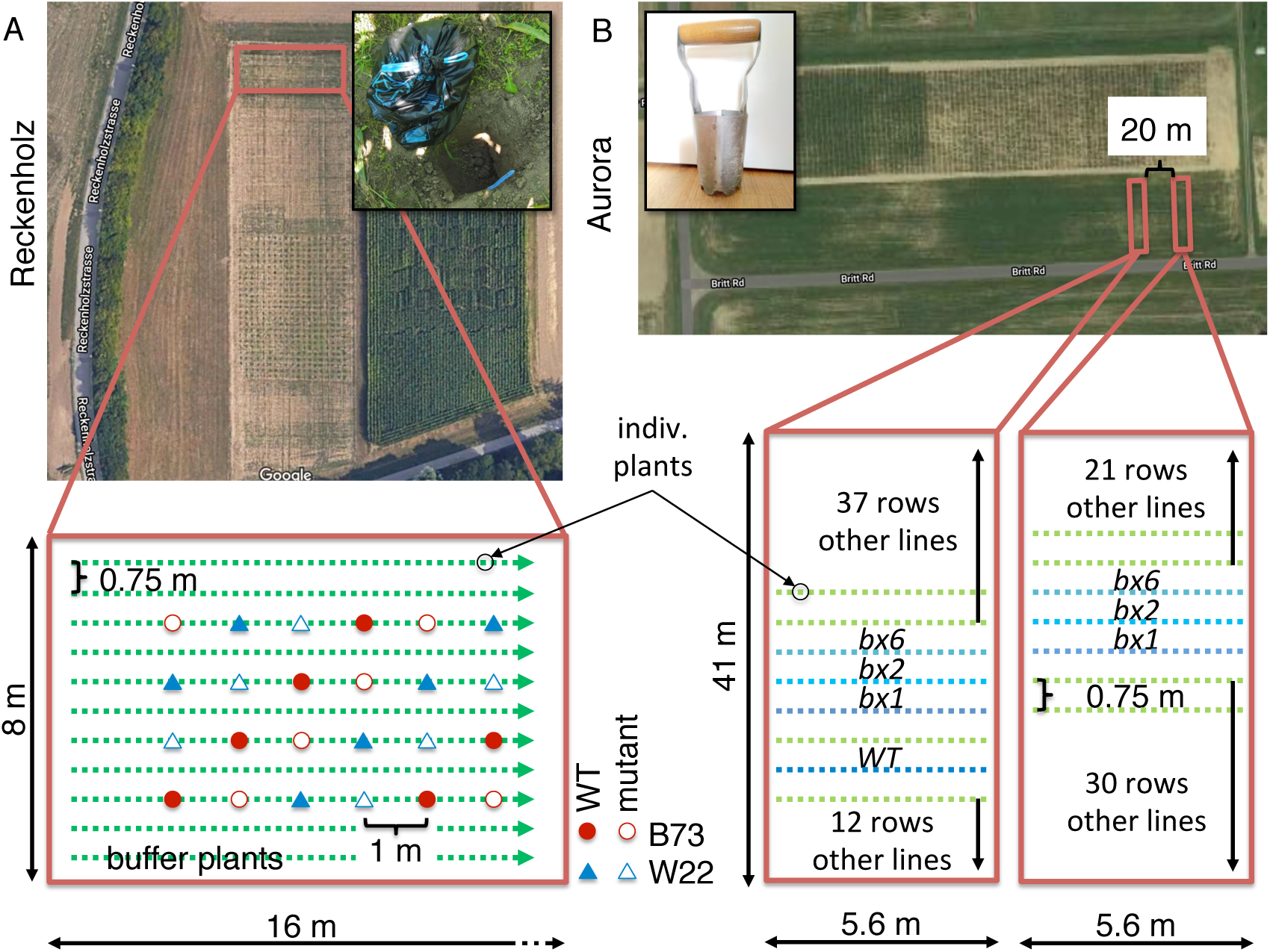
Setup of field experiments The aerial views display the fields used for the experiment in Reckenholz (**A**, Parcel 209) and Aurora (**B**, Field U). The fields can be located with the coordinates given in the methods. Red boxes frame the areas of the fields that were used and they link to a scheme of the experimental designs. In Reckenholz, the test plants were spaced by 1 m between buffer plants whereas the test plants in Aurora were sown in rows. Plant genotypes are indicated by symbols and color (A) or directly labeled (B). The inlets visualize the sampling methods to collect the soil cores (A) or the soil cylinders (B).

**Figure S2.**
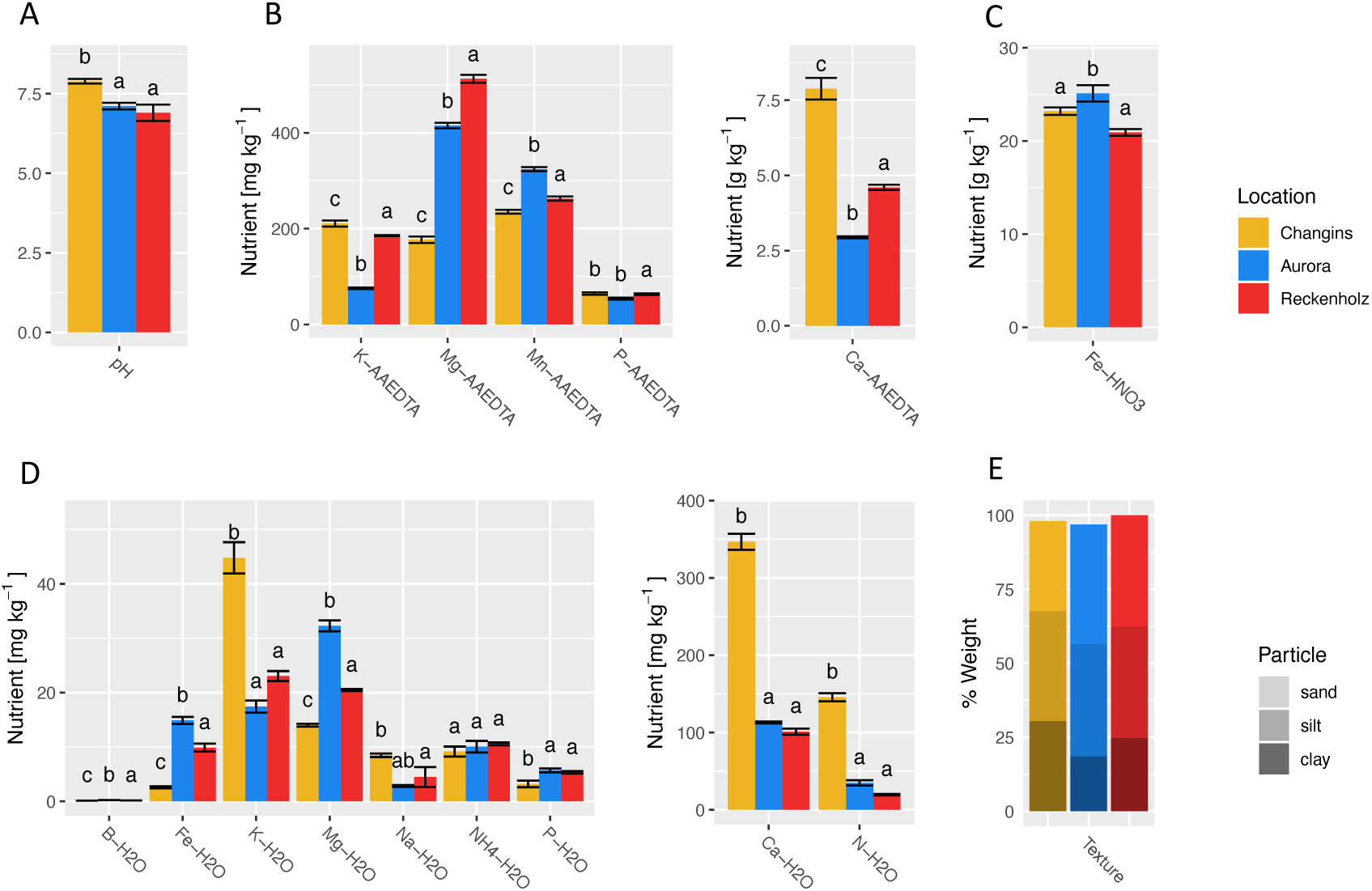
Soil characteristics Physical and chemical characteristics of the three soils from the locations Changins, Aurora and Reckenholz. (A) pH was determined with H2O; soils chemical characteristics were determined in (B) 1:10 acetate-ammonium EDTA (AAEDTA, proxy for reserve nutrients) extracts and (D) 1:10 water (H2O, proxy for plant available nutrients), (C) total iron was measured in nitric acid (HNO3) extracts and (E) soil texture was measured by fractionation. Letters indicate significant differences between locations (*P*-value < 0.05, Tukey HSD test).

**Figure S3.**
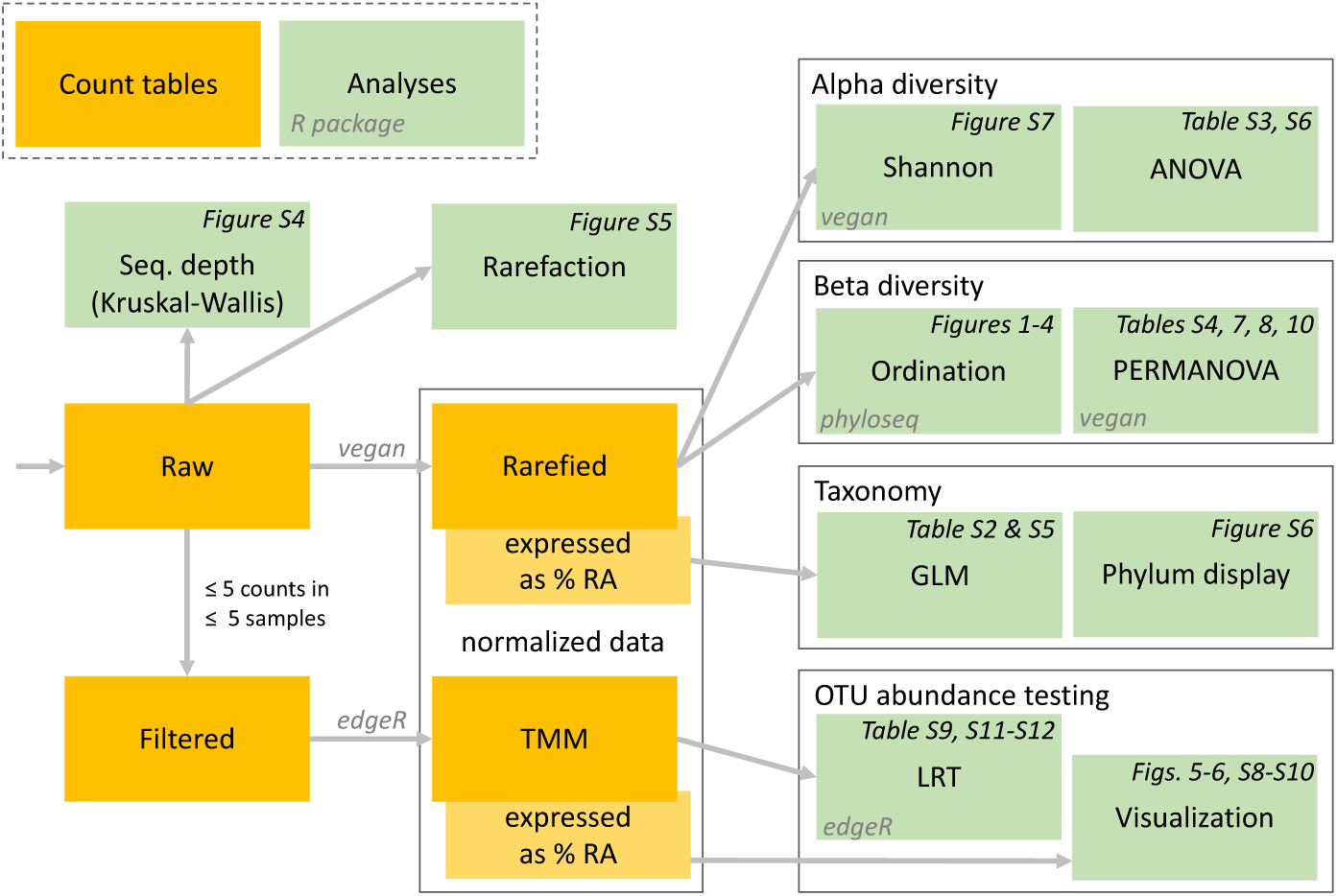
Analysis steps The schematic flow diagram illustrates the steps of the analysis in R. Individual types of normalization steps, analyses or statistical tests are indicated with the blue boxes. Larger grey boxes segment the analysis and indicate the major R-packages that were used in alpha- and beta diversity analyses, differential abundance testing and network analysis. Analysis outputs (Figures and Tables) are indicated in red at their respective analysis steps.

**Figure S4.**
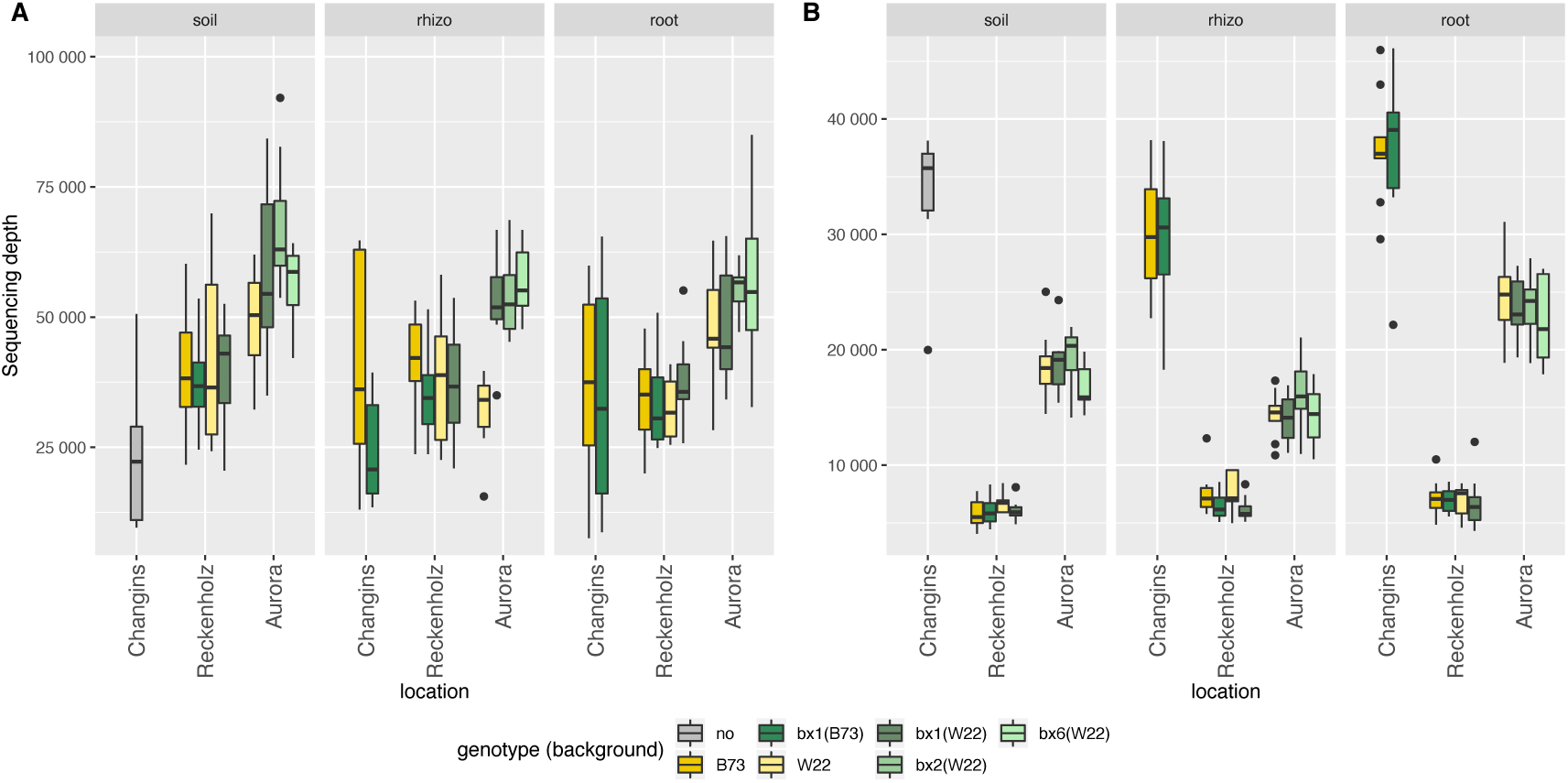
Sequencing effort by sample groups Sequencing depths of (A) bacterial and (B) fungal community profiles. Panels show different compartments, and the different genotypes for each location are shown within the panels. Mean sequencing depths differ significantly between groups of samples (Kruskal-Wallis test, *P* < 0.05, **Supplementary Data S3**).

**Figure S5.**
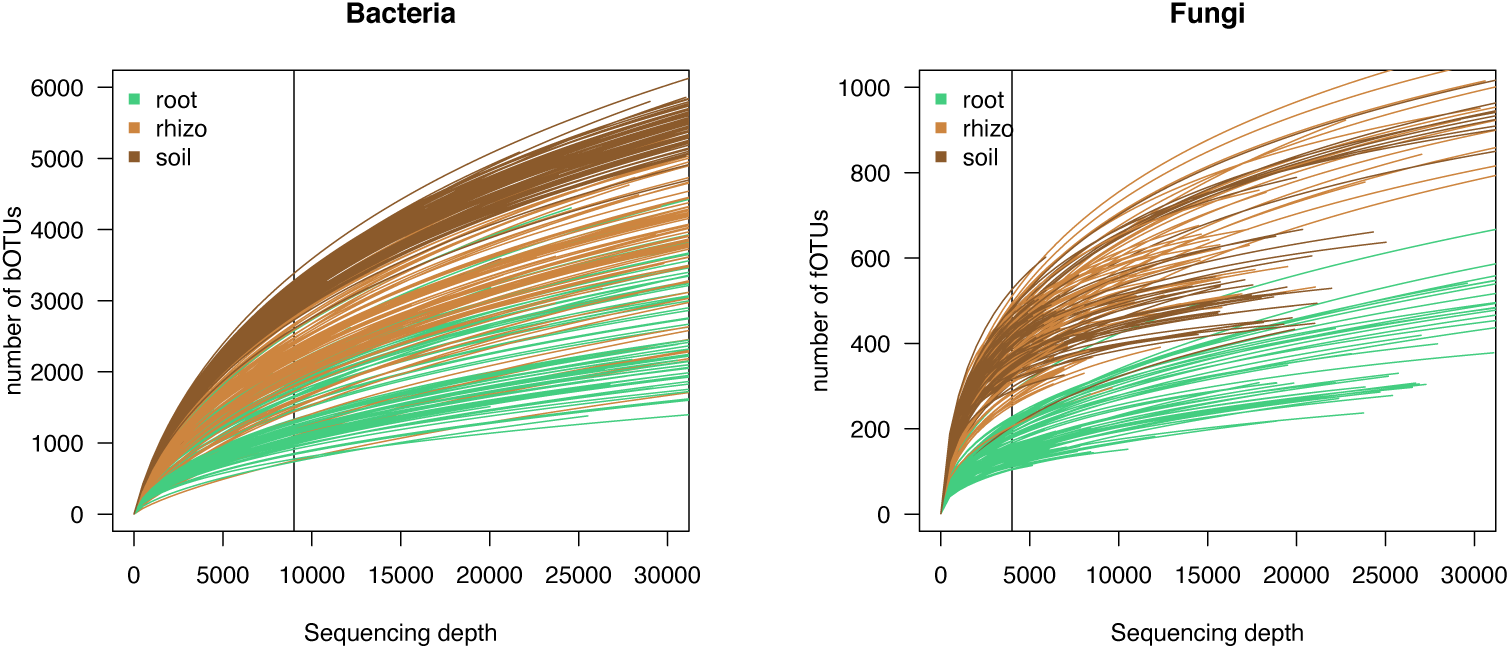
Sampling intensity analysis Rarefaction curve for (A) bacterial and (B) fungal OTU richness. Black lines indicate rarefaction thresholds (9,000 and 4,000 sequences per sample) used for alpha diversity analysis.

**Figure S6.**
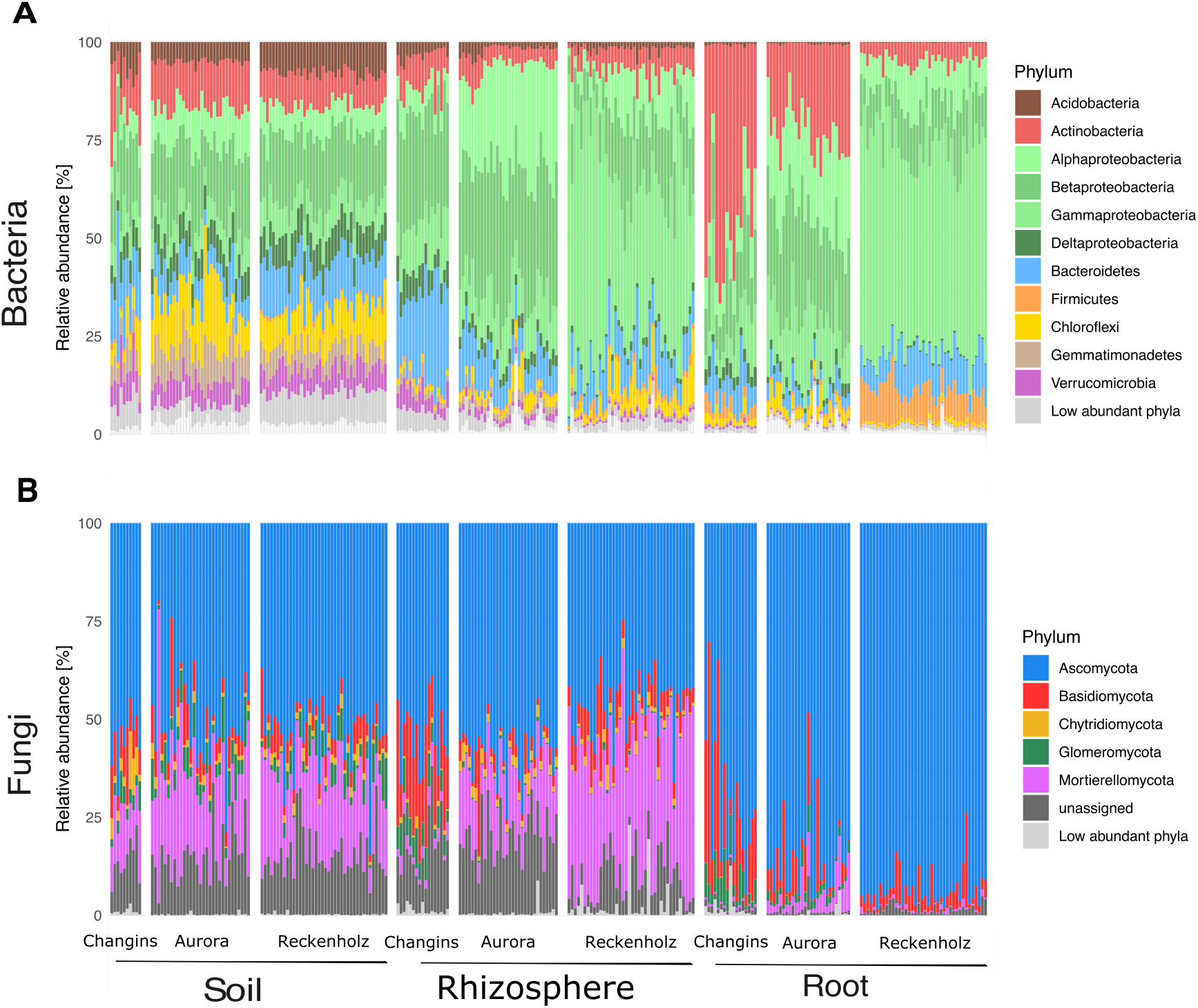
Taxonomy Taxonomy profiles of (A) bacteria (B) fungi at the class level for Proteobacteria and at the phylum level for all other phyla. Different facets in one plot show different samples (sample compartment and location), with all genotypes and replicates. Phyla are considered low abundant when they are below 1% of relative abundance. The statistical testing between the different locations and compartments is documented in **Table S2** and the analysis of BX effects in each compartment in **Table S5**.

**Figure S7.**
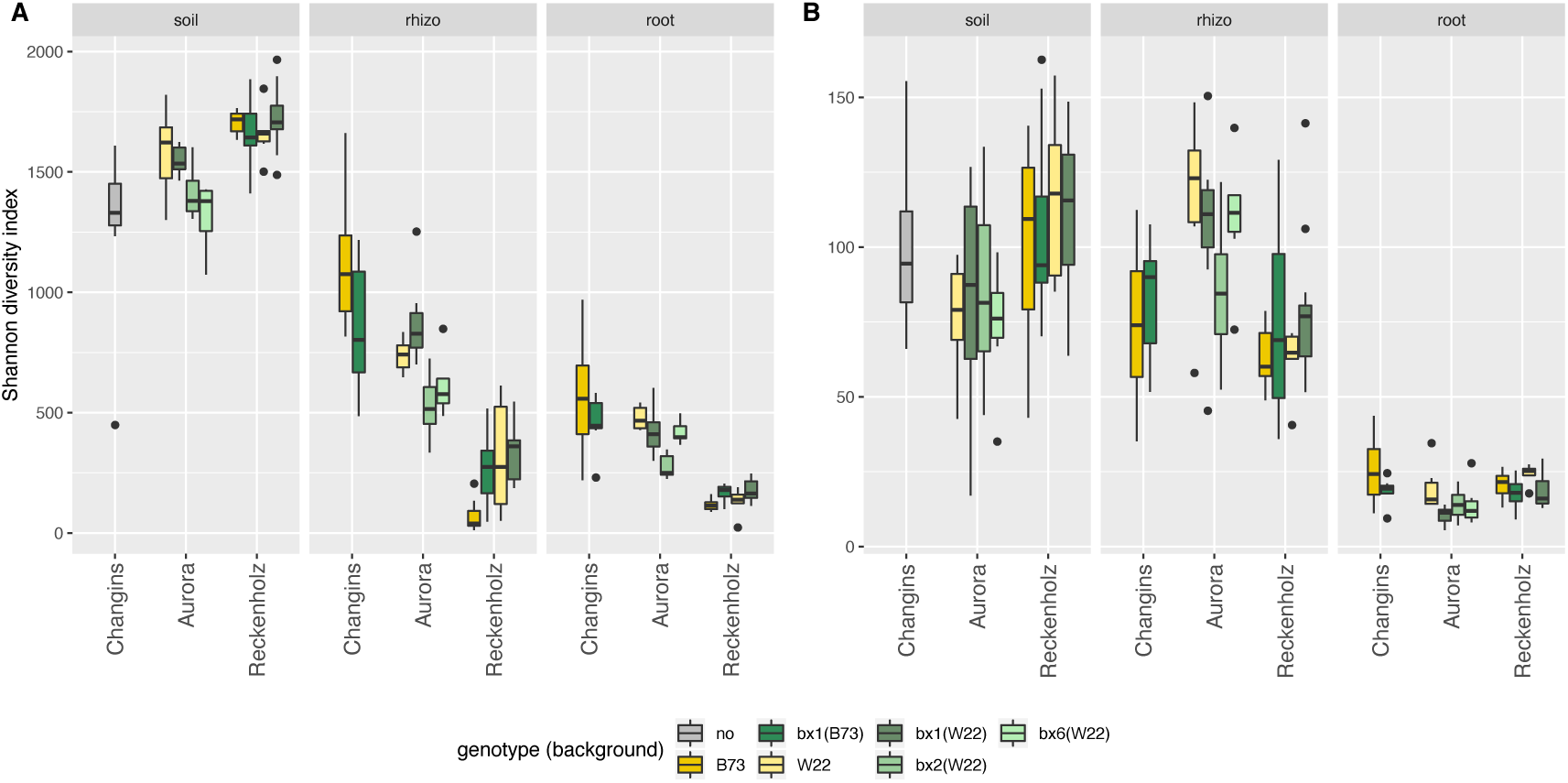
Alpha diversity Alpha diversity was measured with the Shannon index for (A) bacteria and (B) fungi on rarefied data (9,000 and 4,000 sequences per sample for bacteria and fungi, respectively), for different sample compartment (plot facets) and locations. The statistical testing between the different locations and compartments is documented in **Table S3**.

**Figure S8.**
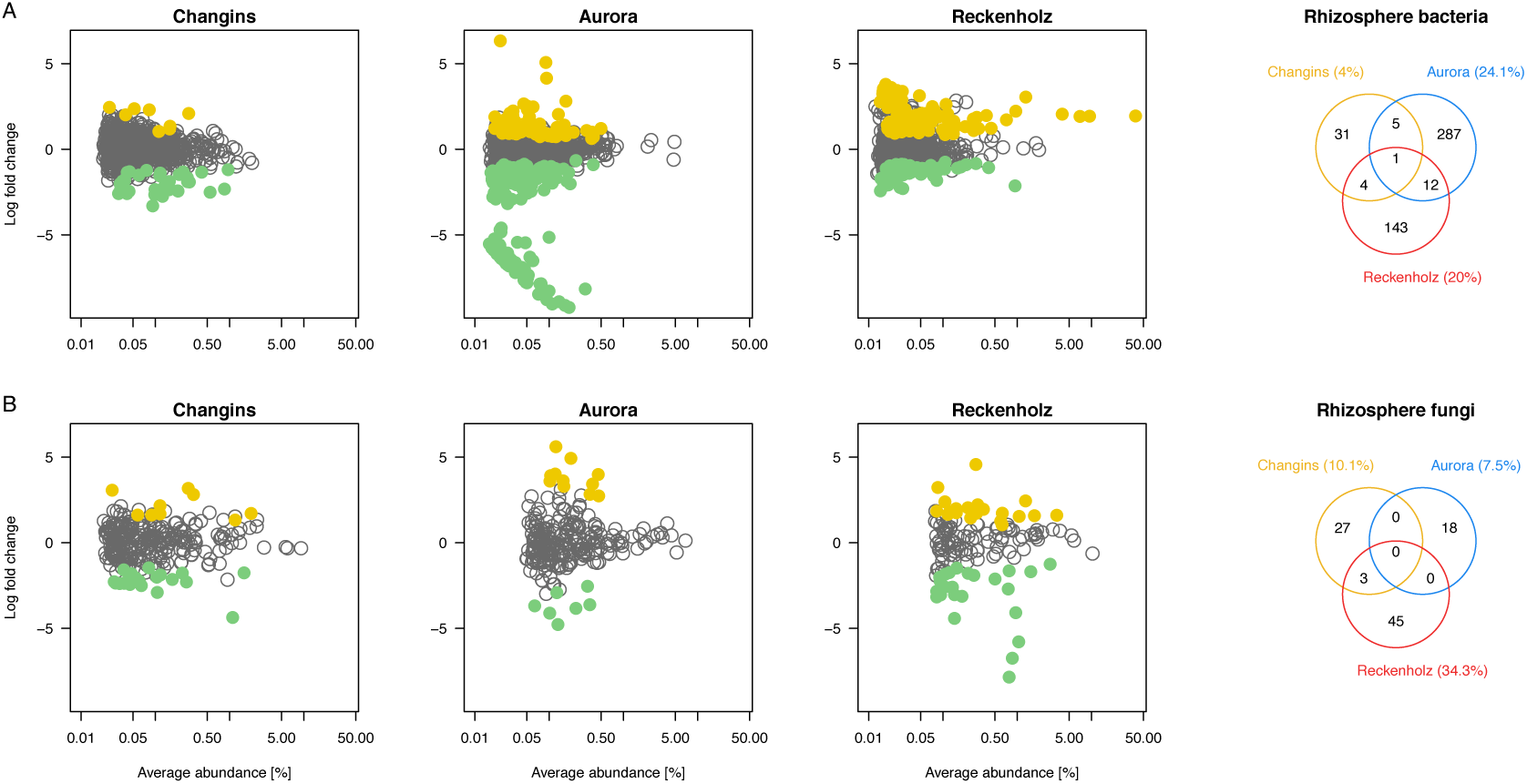
BX-sensitive rhizosphere microbes across locations The MA plots display the average abundance (in log count per million, CPM) and the log-fold change of all b/fOTUs plotted on the x- and y-axes, respectively. b/fOTUs being differentially abundant between wild-type and *bx1* mutant lines (BX-sensitive OTUs) were determined by edgeR analysis (FDR < 0.05, **Table S12**). Colors refer to enriched b/fOTUs in wild-type (yellow) or *bx1* mutant (green) lines. (A) reports the rhizosphere bacteria and (B) the rhizosphere fungi at the locations Changins (yellow), Aurora (blue) and Reckenholz (red). The comparison of BX-sensitive rhizosphere b/fOTUs between locations is visualized with the Venn diagrams.

**Figure S9.**
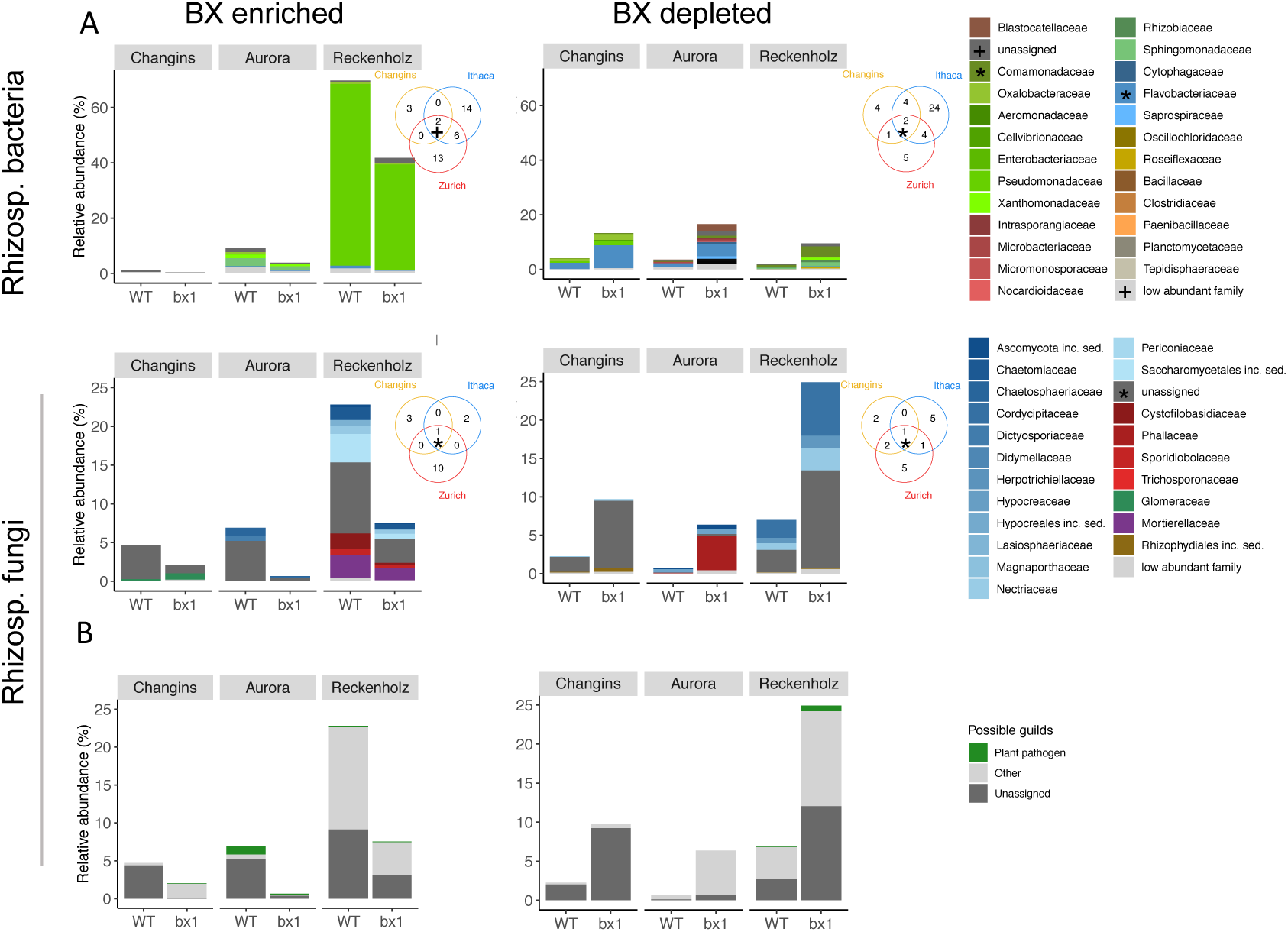
Taxonomic patterns of BX-sensitive rhizosphere microbes The bar plots depict for each location the mean relative abundances (in %) and taxonomies of all rhizosphere bOTUs (upper panels) and rhizosphere fOTUs (lower panels) that differed significantly in abundance between wild-type (WT) and *bx1* mutant lines (i.e., the BX-sensitive b/fOTUs as determined by edgeR analysis, FDR < 0.05, **Table S12**). The BX-enriched (left panels) and BX-depleted taxa (right panels) correspond to the same yellow (enriched in WT) and green (enriched in *bx1*) b/fOTUs of **Fig. S8**, respectively. Individual b/fOTUs are displayed in a stacked manner sorted by their taxonomic assignment at family level. The Venn diagram insets compare the family assignments of the BX-sensitive taxa between the locations Changins (yellow), Aurora (blue) and Reckenholz (red). Overlapping family assignments are indicated in the plot or marked in the taxonomy legend. (B) visualizes the proportion of assignments to ‘plant pathogen’ among the FUNGuild annotations. The sets of BX-enriched and BX-depleted rhizosphere fOTUs from each location were annotated individually to their ecological guilds.

**Figure S10.**
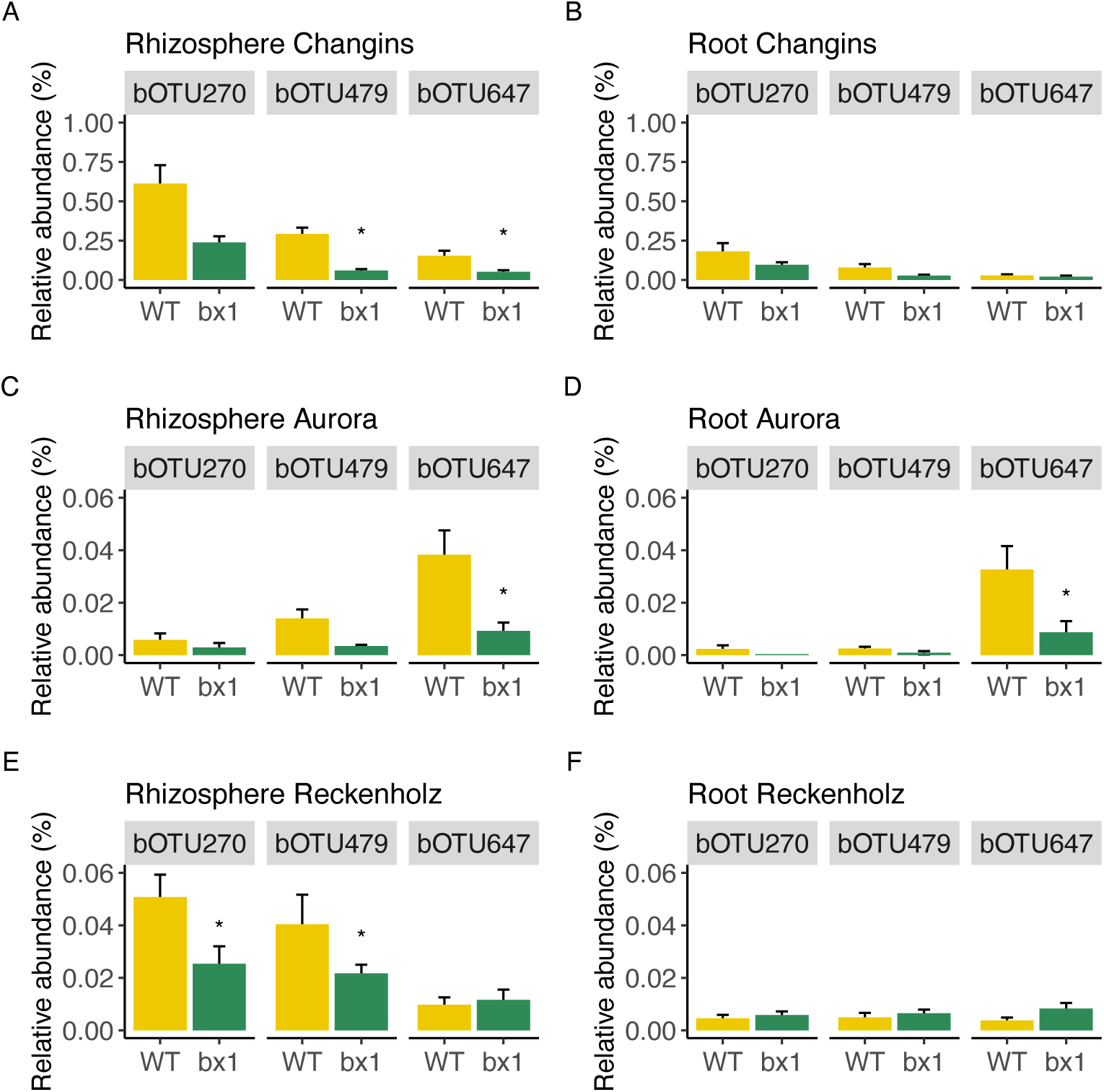
Abundance of *Methylophilaceae* bOTUs across compartments and locations Bar graphs display the mean abundance (± s.e.m.) of the *Methylophilales* bOTU479 and the two *Methylophilaceae* bOTUs #270 and #647 in rhizosphere (A,C,E) and root (B,D,F) samples of wild-type (WT) and *bx1* mutant lines at all three locations. Asterisks mark significant differences between WT and *bx1* as determined by edgeR analysis (FDR < 0.05, **Table S12**).

## SUPPLEMENTARY TABLES

**Table S1.**
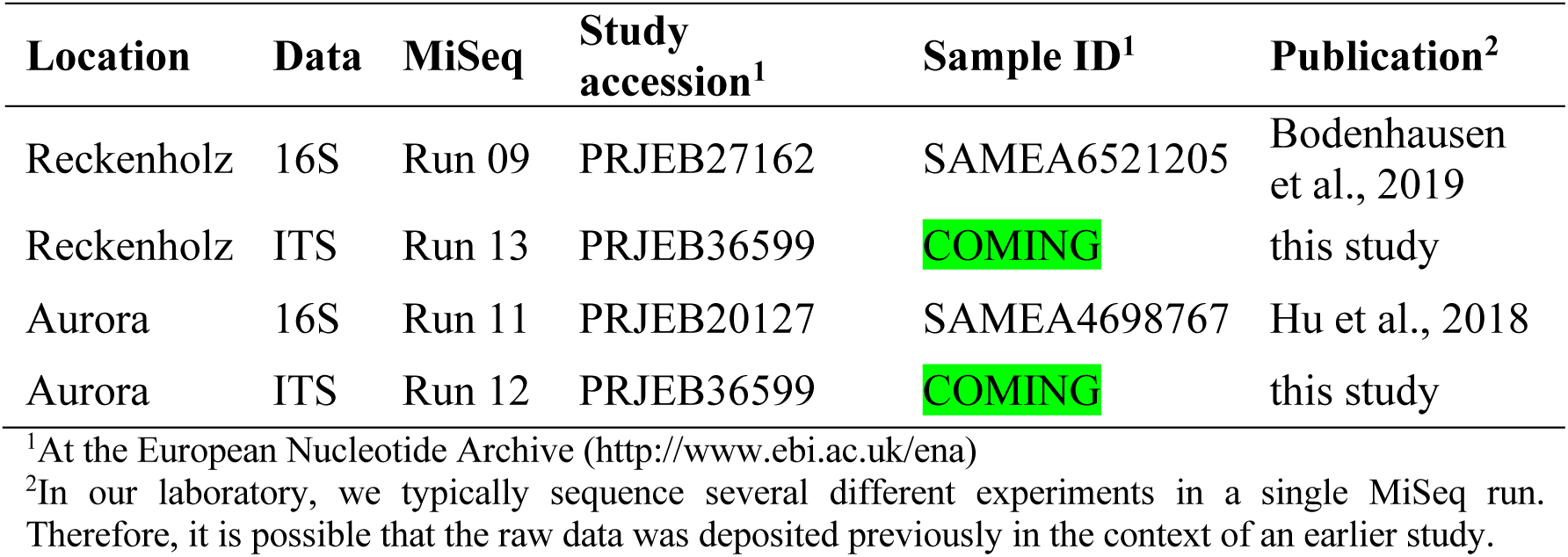
**Deposition of raw sequence data**

**Table S2** Taxonomic analysis comparing locations and compartments The table reports for all sample groups (location * compartment) the F statistic together with the corresponding P value (adjusted for multiple hypothesis testing, following Bonferroni Hochberg) and the relative average abundances [%]. Locations are abbreviated with Ch=Changins, Au=Aurora and Re=Reckenholz. Compartments are abbreviated with So=Soil, Rh=Rhizosphere and Ro=Root. The columns with the posthoc Tukey test results is indicated with “_T” for each sample group. Different letters are used for groups that differ significantly in their abundances (alpha=0.01). -> See supplementary excel file (Table_S2_Phylum_statistics.xlsx)

**Table S3.**
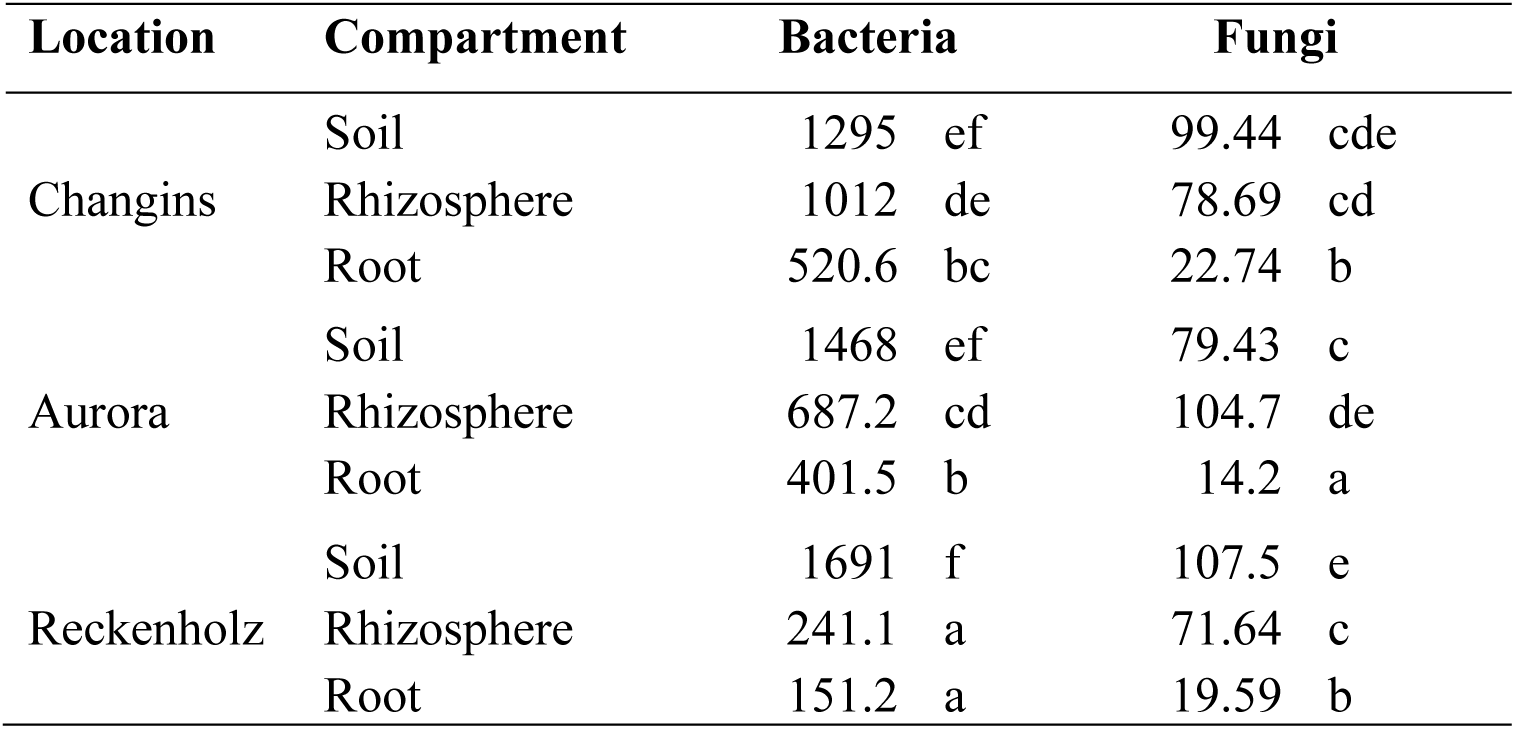
**Alpha diversity analysis comparing locations and compartments** Statistics of Shannon diversity between compartments (soil, rhizosphere and root) at each location (Changins, Aurora and Reckenholz; Tukey HSD test). Different letters correspond to groups that are significantly different from each other in Shannon diversity at *P* < 0.05.

**Table S4.**
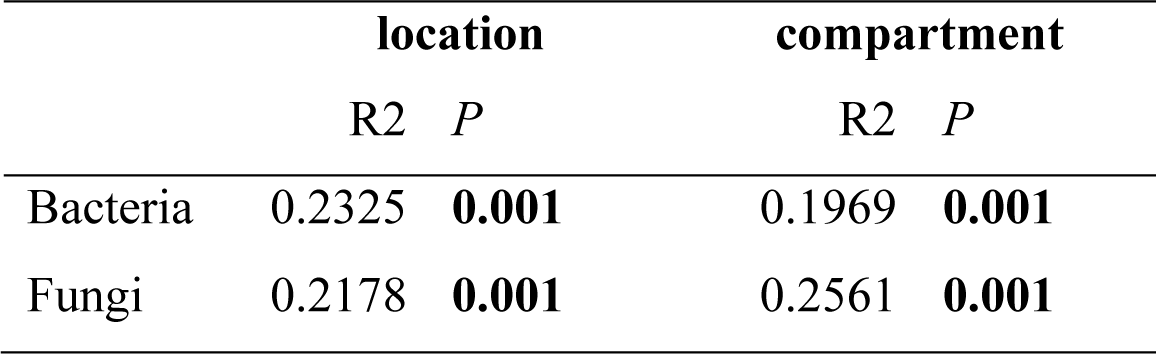
Beta diversity analysis comparing locations and compartments Effects of location and compartment on beta diversity of bacterial and fungal diversity. R2 and *P*-values of PERMANOVA on Bray-Curtis distances are presented.

**Table S5** Taxonomic analysis comparing BX effects in each compartment The table reports for bacteria and fungi and for each compartment the F statistic together with the corresponding P value (adjusted for multiple hypothesis testing, following Bonferroni Hochberg). The table also displays the relative average abundances [%] per sample group (location * genotype). Locations are abbreviated with Ch=Changins, Au=Aurora and Re=Reckenholz. Genotypes are abbreviated with WT=wildtype and bx for the bx1 mutant. The columns with the posthoc Tukey test results is indicated with “_T” for each sample group. Different letters are used for groups that differ significantly in their abundances (alpha=0.01). -> See supplementary excel file (Table_S5_Phylum_BX_statistics.xlsx)

**Table S6.**
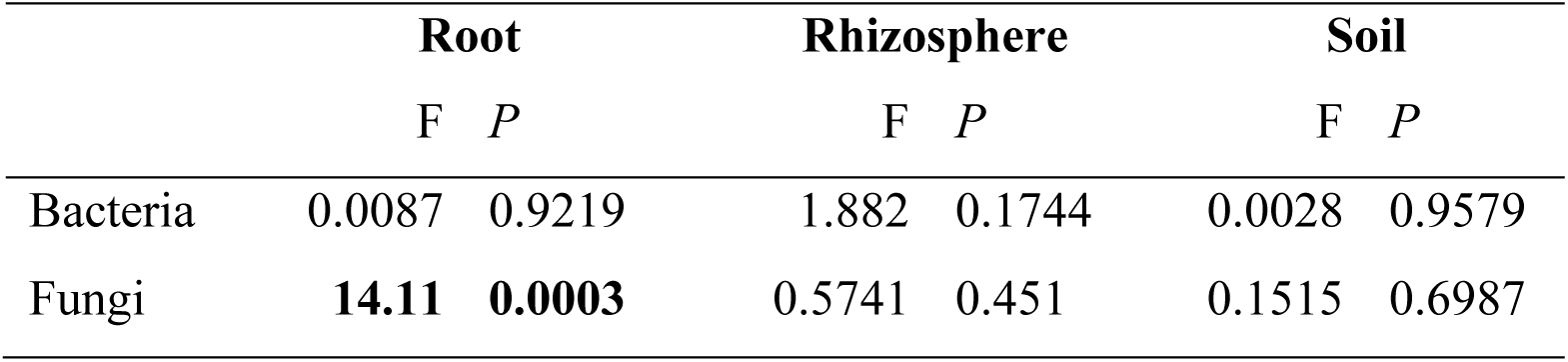
**Alpha diversity analysis comparing BX effects in each compartment** Effect of BX production (Wild type vs. mutant) on Shannon diversity within compartments. *F* and *P*-values (if significant, then in bold) from ANOVA are presented.

**Table S7.**
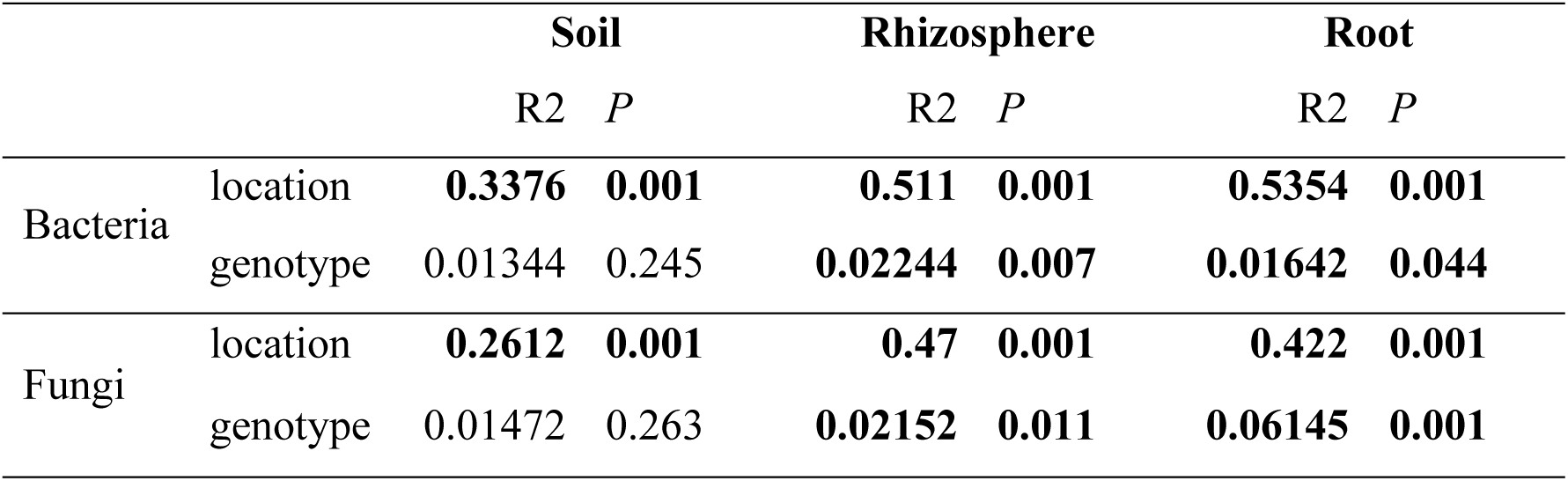
Beta diversity analysis comparing BX and location effects in each compartment Location and genotype effects of BX-exudation on beta diversity of bacterial and fungal communities within compartments. R2 and *P*-values of PERMANOVA on Bray-Curtis distance are presented. Significant effects are in bold.

**Table S8.**
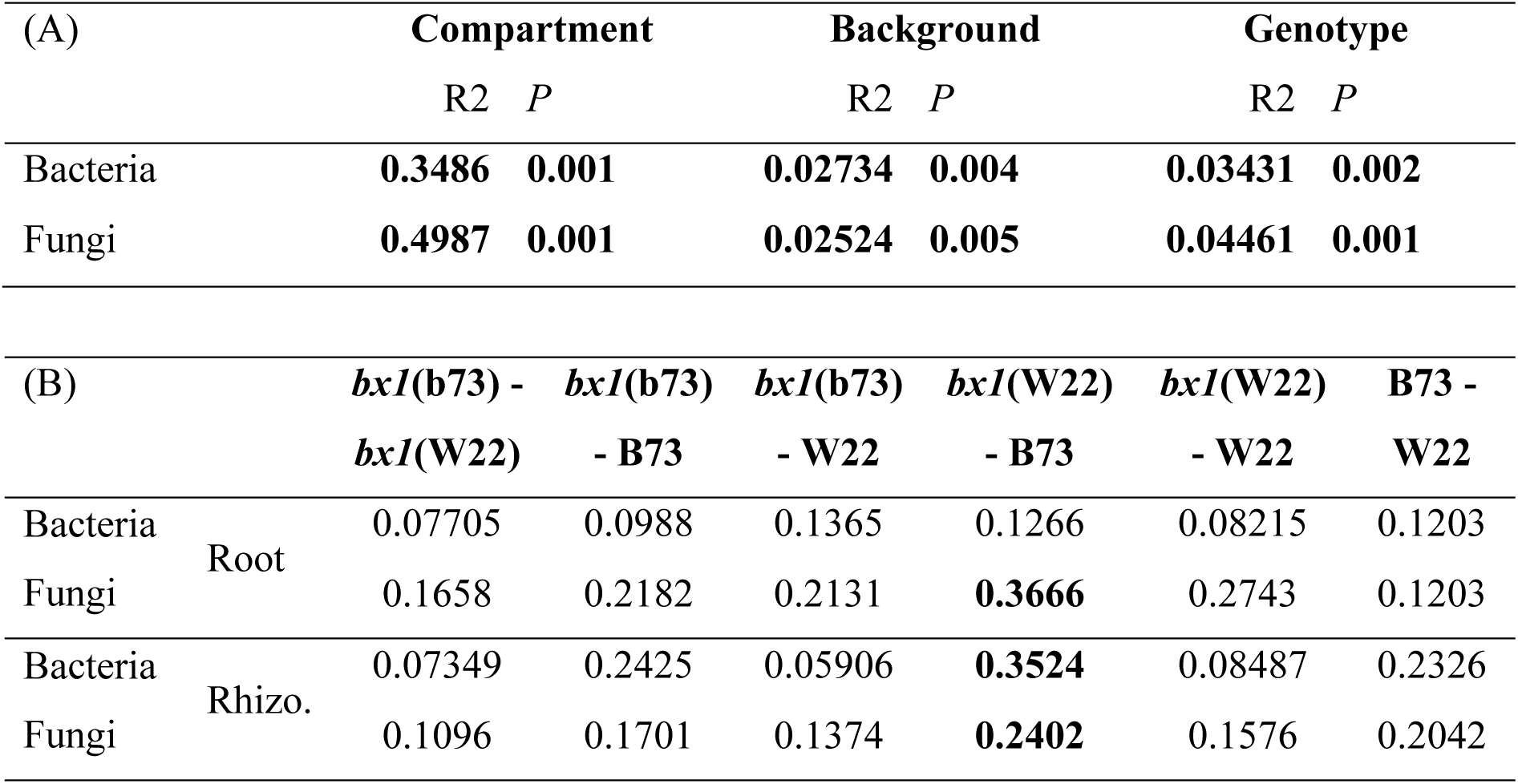
Beta diversity analysis comparing background and genotype effects in each compartment (Reckenholz experiment) Factorial (A) PERMANOVA analysis and (B) pairwise PERMANOVA analysis of bacterial and fungal communities at the Reckenholz location testing for effects of compartment (soil, root and rhizosphere), genetic background (B73, W22) and plant genotype (WT, *bx1*). R2 and *P*-values of PERMANOVA on Bray-Curtis distance are presented. For pairwise PERMANOVA, R2 values are reported and significant (FDR < 0.05) effects marked in bold.

**Table S9** zOTUs differing by background or genotype (Reckenholz experiment) The supplementary table lists, for bacteria and fungi of the root and the rhizosphere, all zOTUs differing significantly between backgrounds (B73, W22) or genotypes (WT, *bx1*) based on edgeR analysis. The table lists the zOTU-ID, taxonomy, the mean log abundance in counts per million [logCPM] and log fold change [logFC] between backgrounds or genotypes and the statistic of the likelihood ratio test [LR], its *P*-value [PValue] and its FDR-corrected *P*-value [FDR]. -> See supplementary excel file (Table_S9_Reckenholz_OTU_statistics.xlsx)

**Table S10.**
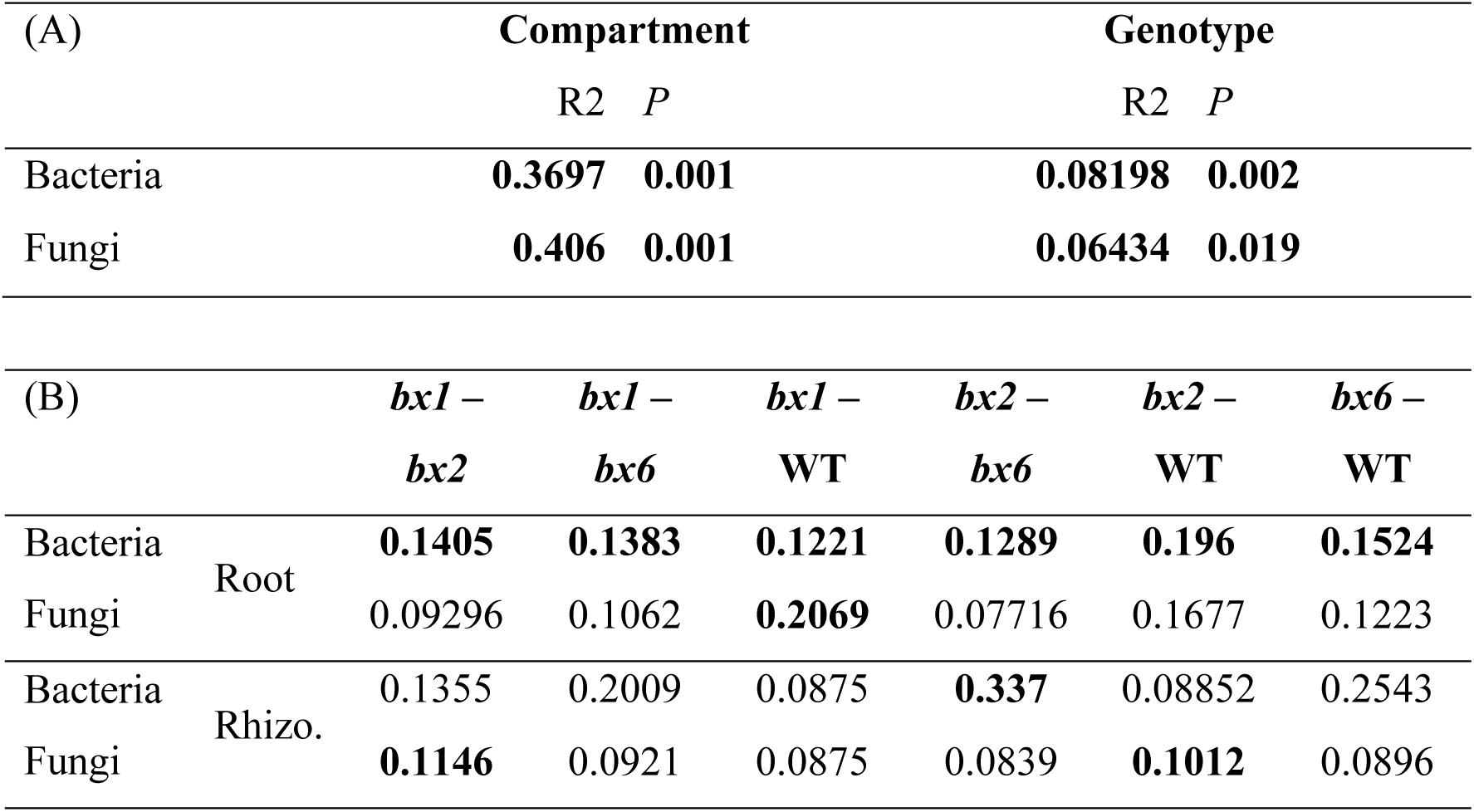
Beta diversity analysis comparing mutant genotypes in each compartment (Aurora experiment) Factorial (A) PERMANOVA analysis and (B) pairwise PERMANOVA analysis of bacterial and fungal communities at the Aurora location testing for effects of compartment (soil, root and rhizosphere) and plant genotypes (WT, *bx1*, *bx2* and *bx6*). R2 and *P*-values of PERMANOVA on Bray-Curtis distance are presented. For pairwise PERMANOVA, R2 values are reported and significant (FDR < 0.05) effects marked in bold.

**Table S11** zOTUs differing by mutant genotypes (Aurora experiment) The supplementary table lists, for bacteria and fungi of the root and the rhizosphere, all zOTUs differing significantly between among WT and the mutant lines based on edgeR analysis. The table lists the zOTU-ID, taxonomy, the mean log abundance in counts per million [logCPM] and log fold change [logFC] between WT and *bx1* and the statistic of the likelihood ratio test [LR], its *P*-value [PValue] and its FDR-corrected *P*-value [FDR]. -> See supplementary excel file (Table_S11_Aurora_OTU_statistics.xlsx)

**Table S12** BX-sensitive zOTUs across all locations The supplementary excel table lists for bacteria and fungi of the root and the rhizosphere all zOTUs differing significantly between WT and *bx1* mutant lines for each location based on edgeR analysis. That table lists the location, the zOTU-ID, taxonomy, the mean log abundance in counts per million [logCPM] and log fold change [logFC] between WT and *bx1* and the statistic of the likelihood ratio test [LR], its *P*-value [PValue] and its FDR-corrected *P*-value [FDR]. -> See supplementary excel file (Table_S12_BX_OTU_statistics.xlsx)

