## Supplementary Information for "Specific and conserved patterns of microbiota-structuring by maize benzoxazinoids in the field"

### Cadot et al. (2020, JOURNAL)

**INDEX:**

---

**SUPPLEMENTARY RESULTS**

**SUPPLEMENTARY FIGURES**

Figure S1 | Setup of field experiments
Figure S2 | Soil characteristics
Figure S3 | Analysis steps
Figure S4 | Sequencing effort by sample groups
Figure S5 | Sampling intensity analysis
Figure S6 | Taxonomy
Figure S7 | Alpha diversity
Figure S8 | BX-sensitive rhizosphere microbes across locations
Figure S9 | Taxonomic patterns of BX-sensitive rhizosphere microbes
Figure S10 | Abundance of Methylophilaceae bOTUs across compartments and locations

**SUPPLEMENTARY TABLES:**

Table S1 | Deposition of raw sequence data
Table S2 | Taxonomic analysis comparing locations and compartments
Table S3 | Alpha diversity analysis comparing locations and compartments
Table S4 | Beta diversity analysis comparing locations and compartments
Table S5 | Taxonomic analysis comparing BX effects in each compartment
Table S6 | Alpha diversity analysis comparing BX effects in each compartment
Table S7 | Beta diversity analysis comparing BX and location effects in each compartment
Table S8 | Beta diversity analysis comparing background and genotype effects in each compartment (Reckenholz experiment)
Table S9 | zOTUs differing by background or genotype (Reckenholz experiment)
Table S10 | Beta diversity analysis comparing mutant genotypes in each compartment (Aurora experiment)
Table S11 | zOTUs differing by mutant genotypes (Aurora experiment)
Table S12 | BX-sensitive zOTUs across all locations

### SUPPLEMENTARY FIGURES:

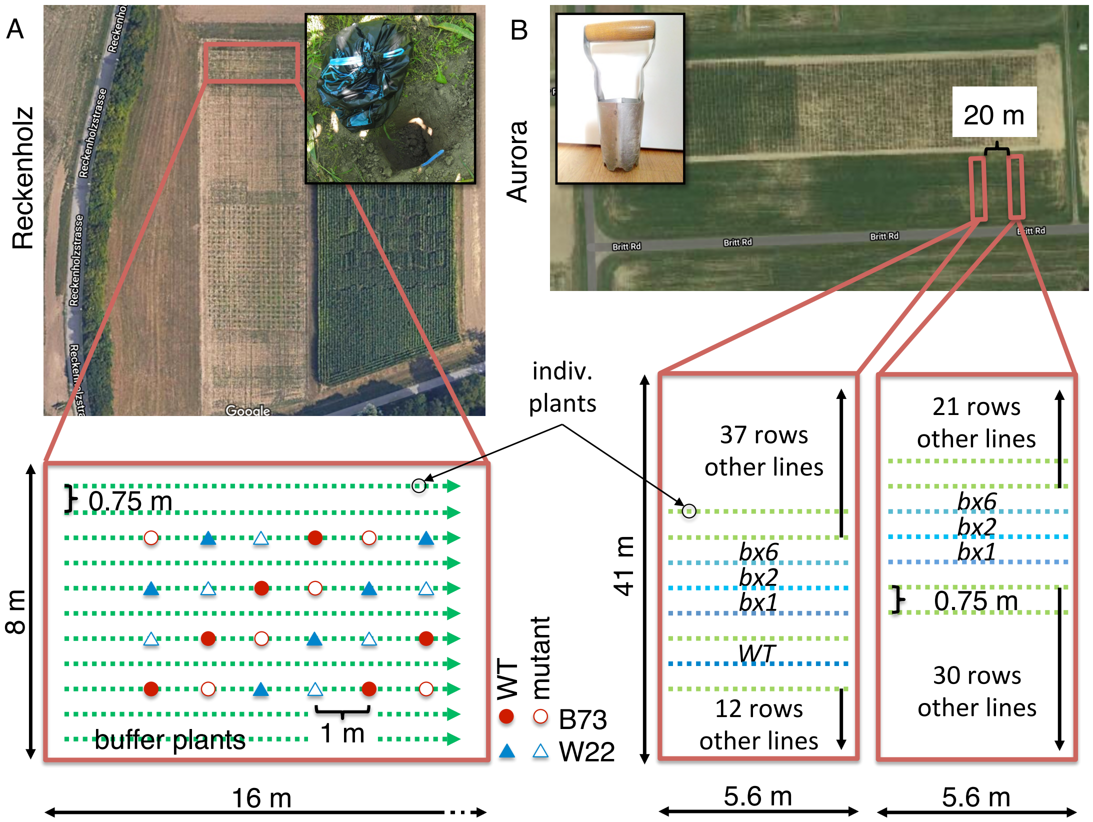


**Figure S1 | Setup of field experiments**

The aerial views display the fields used for the experiment in Reckenholz (**A**, Parcel 209) and Aurora (**B**, Field U). The fields can be located with the coordinates given in the methods. Red boxes frame the areas of the fields that were used and they link to a scheme of the experimental designs. In Reckenholz, the test plants were spaced by 1 m between buffer plants whereas the test plants in Aurora were sown in rows. Plant genotypes are indicated by symbols and color (A) or directly labeled (B). The inlets visualize the sampling methods to collect the soil cores (A) or the soil cylinders (B).

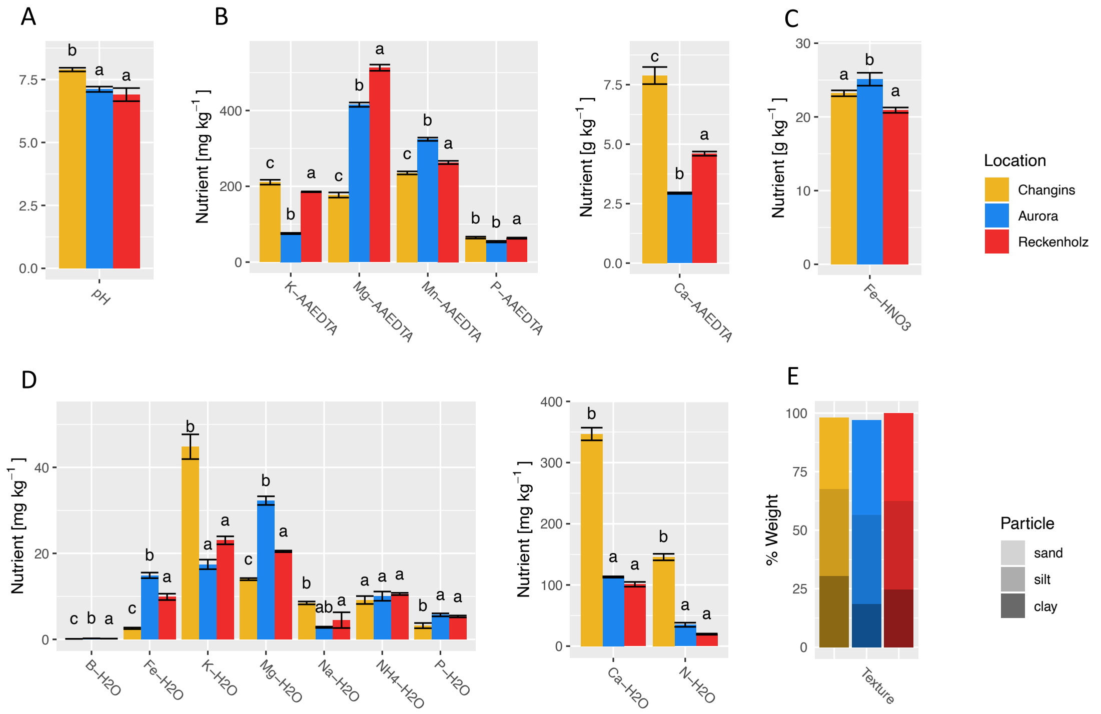


**Figure S2 | Soil characteristics**

Physical and chemical characteristics of the three soils from the locations Changins, Aurora and Reckenholz. (A) pH was determined with $H_2O$; soils chemical characteristics were determined in (B) 1:10 acetate-ammonium EDTA (AAEDTA, proxy for reserve nutrients) extracts and (D) 1:10 water (H2O, proxy for plant available nutrients), (C) total iron was measured in nitric acid (HNO3) extracts and (E) soil texture was measured by fractionation. Letters indicate significant differences between locations (*P*-value < 0.05, Tukey HSD test).

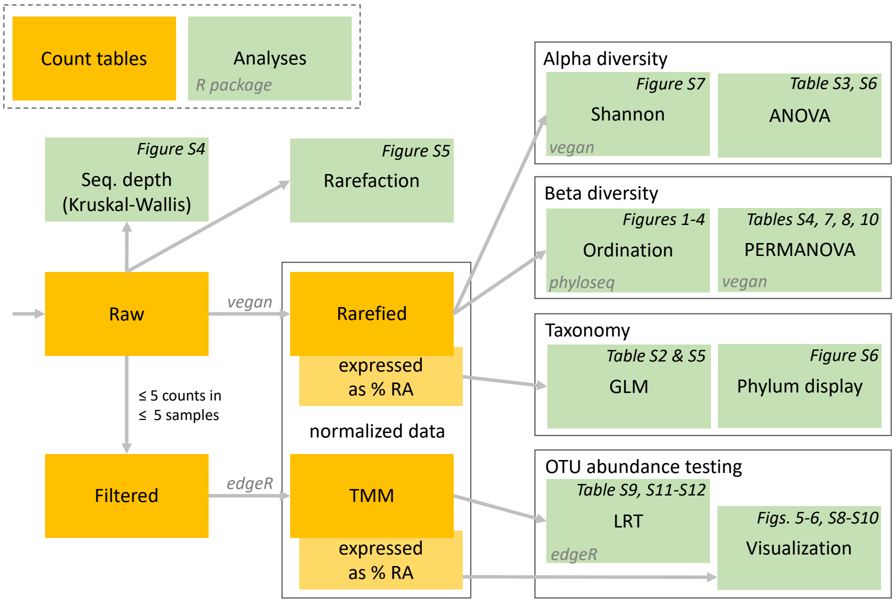


**Figure S3 | Analysis steps**

The schematic flow diagram illustrates the steps of the analysis in R. Individual types of normalization steps, analyses or statistical tests are indicated with the blue boxes. Larger grey boxes segment the analysis and indicate the major R-packages that were used in alpha- and beta diversity analyses, differential abundance testing and network analysis. Analysis outputs (Figures and Tables) are indicated in red at their respective analysis steps.

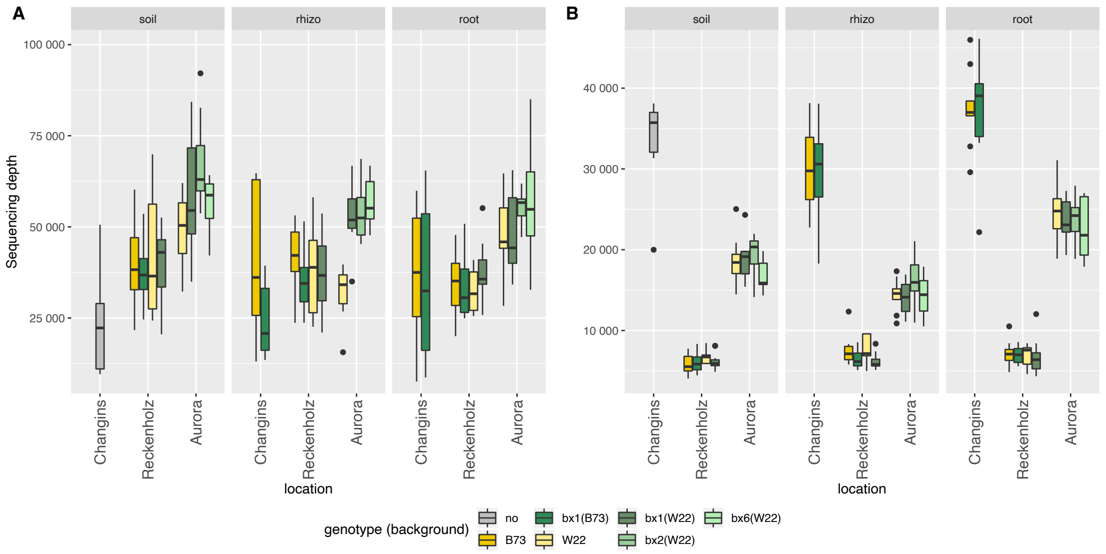


**Figure S4 | Sequencing effort by sample groups**

Sequencing depths of (A) bacterial and (B) fungal community profiles. Panels show different compartments, and the different genotypes for each location are shown within the panels. Mean sequencing depths differ significantly between groups of samples (Kruskal-Wallis test, $P < 0.05$, **Supplementary Data S3**).

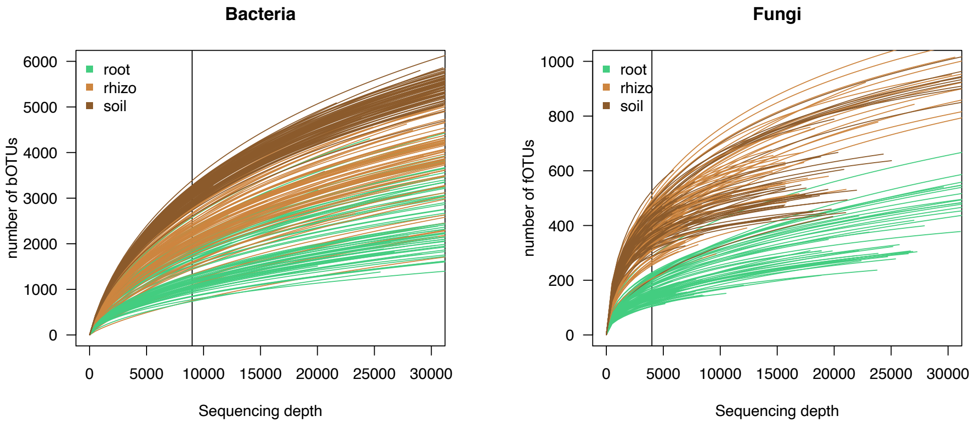


**Figure S5 | Sampling intensity analysis**

Rarefaction curve for (A) bacterial and (B) fungal OTU richness. Black lines indicate rarefaction thresholds (9,000 and 4,000 sequences per sample) used for alpha diversity analysis.

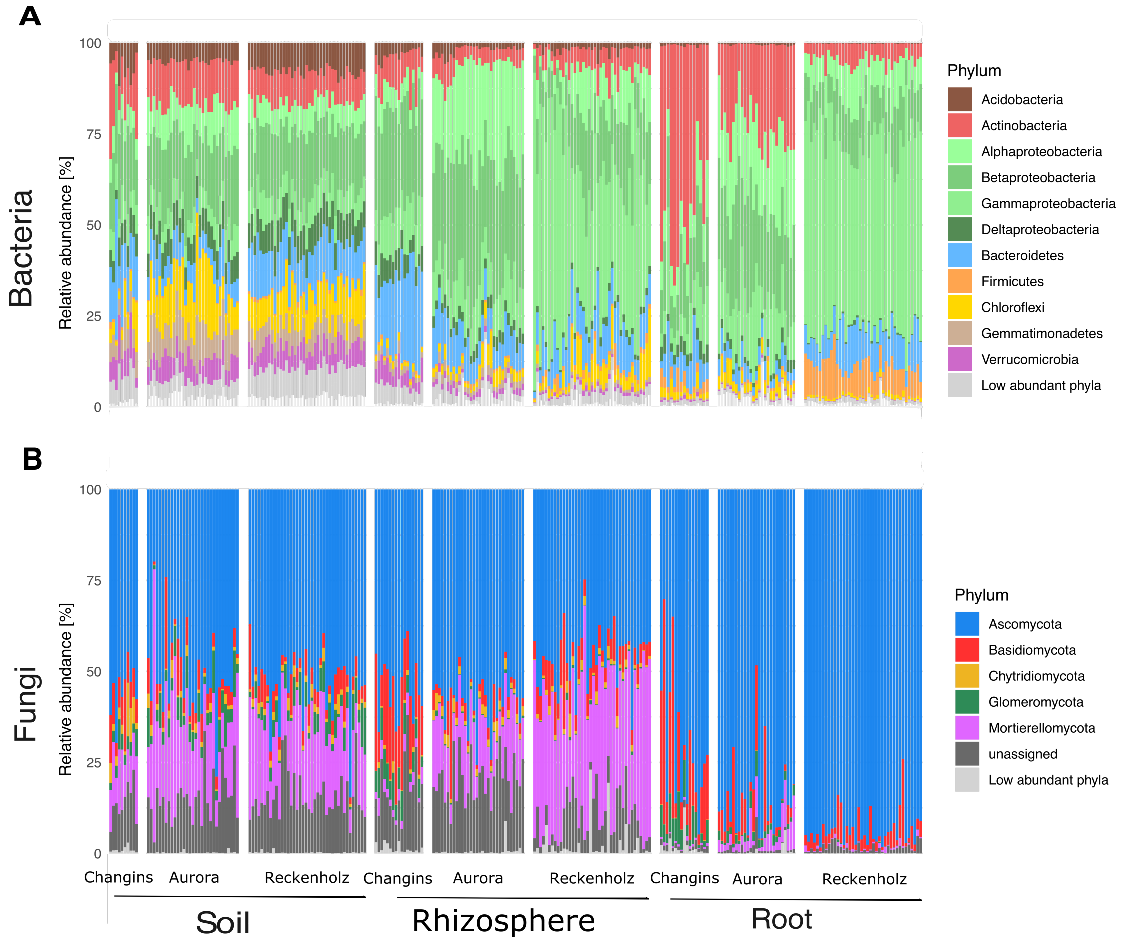


**Figure S6 | Taxonomy**

Taxonomy profiles of (A) bacteria (B) fungi at the class level for Proteobacteria and at the phylum level for all other phyla. Different facets in one plot show different samples (sample compartment and location), with all genotypes and replicates. Phyla are considered low abundant when they are below 1% of relative abundance. The statistical testing between the different locations and compartments is documented in **Table S2** and the analysis of BX effects in each compartment in **Table S5**.

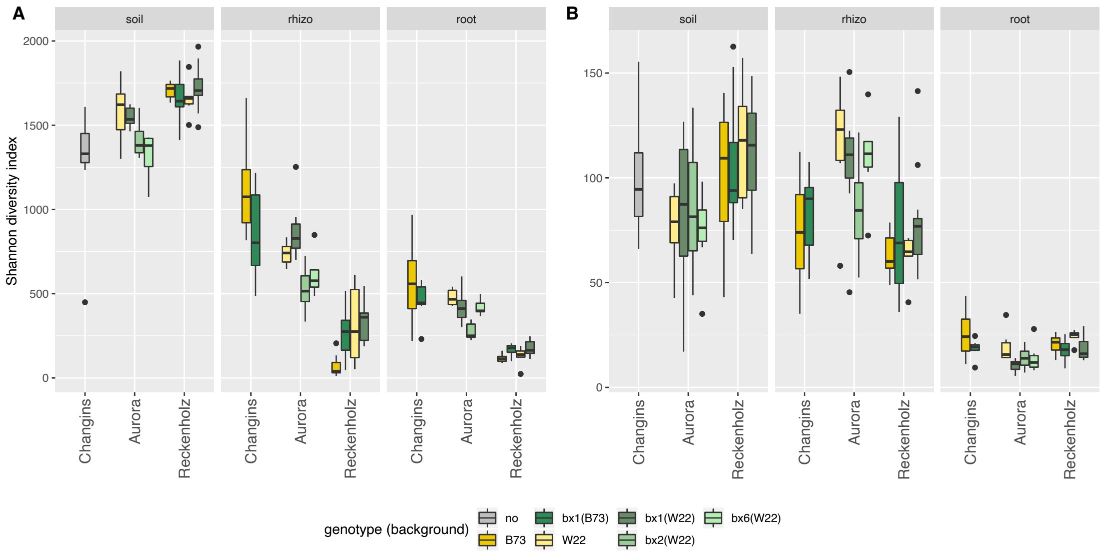


**Figure S7 | Alpha diversity**

Alpha diversity was measured with the Shannon index for (A) bacteria and (B) fungi on rarefied data (9,000 and 4,000 sequences per sample for bacteria and fungi, respectively), for different sample compartment (plot facets) and locations. The statistical testing between the different locations and compartments is documented in **Table S3**.

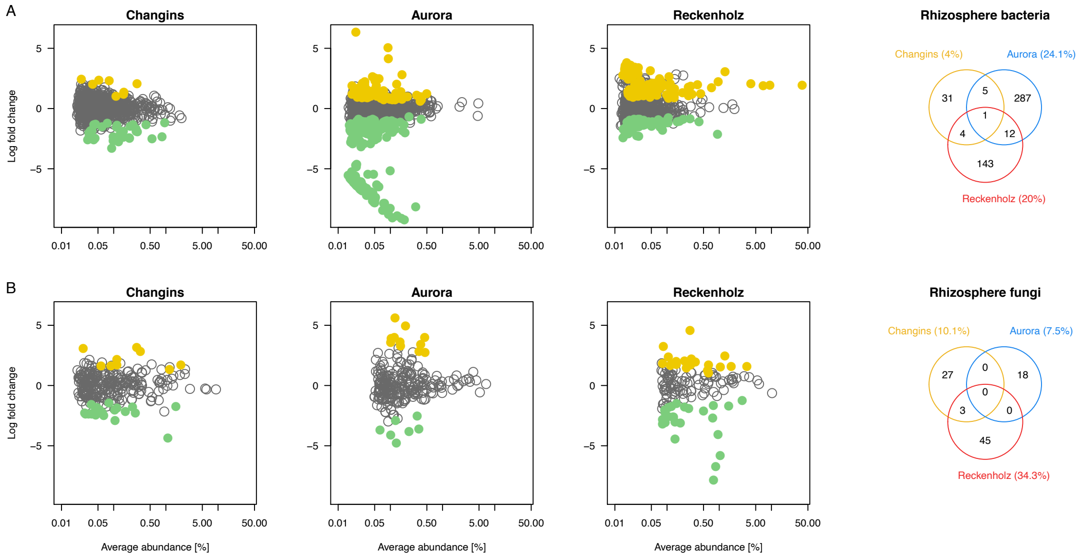


**Figure S8 | BX-sensitive rhizosphere microbes across locations**

The MA plots display the average abundance (in log count per million, CPM) and the log-fold change of all b/fOTUs plotted on the x- and y-axes, respectively. b/fOTUs being differentially abundant between wild-type and *bx1* mutant lines (BX-sensitive OTUs) were determined by edgeR analysis (FDR < 0.05, **Table S12**). Colors refer to enriched b/fOTUs in wild-type (yellow) or *bx1* mutant (green) lines. (A) reports the rhizosphere bacteria and (B) the rhizosphere fungi at the locations Changins (yellow), Aurora (blue) and Reckenholz (red). The comparison of BX-sensitive rhizosphere b/fOTUs between locations is visualized with the Venn diagrams.

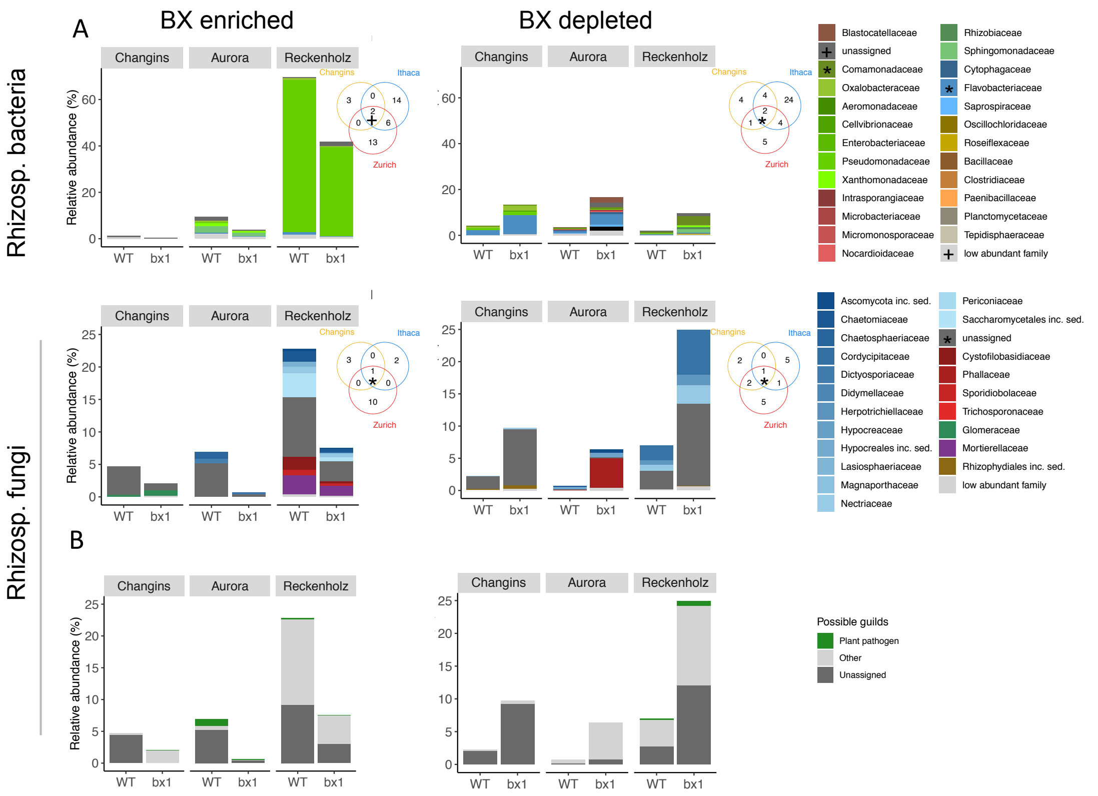


**Figure S9 | Taxonomic patterns of BX-sensitive rhizosphere microbes**

The bar plots depict for each location the mean relative abundances (in %) and taxonomies of all rhizosphere bOTUs (upper panels) and rhizosphere fOTUs (lower panels) that differed significantly in abundance between wild-type (WT) and *bx1* mutant lines (i.e., the BX-sensitive b/fOTUs as determined by edgeR analysis, FDR < 0.05, **Table S12**). The BX-enriched (left panels) and BX-depleted taxa (right panels) correspond to the same yellow (enriched in WT) and green (enriched in *bx1*) b/fOTUs of **Fig. S8**, respectively. Individual b/fOTUs are displayed in a stacked manner sorted by their taxonomic assignment at family level. The Venn diagram insets compare the family assignments of the BX-sensitive taxa between the locations Changins (yellow), Aurora (blue) and Reckenholz (red). Overlapping family assignments are indicated in the plot or marked in the taxonomy legend. (B) visualizes the proportion of assignments to 'plant pathogen' among the FUNGuild annotations. The sets of BX-enriched and BX-depleted rhizosphere fOTUs from each location were annotated individually to their ecological guilds.

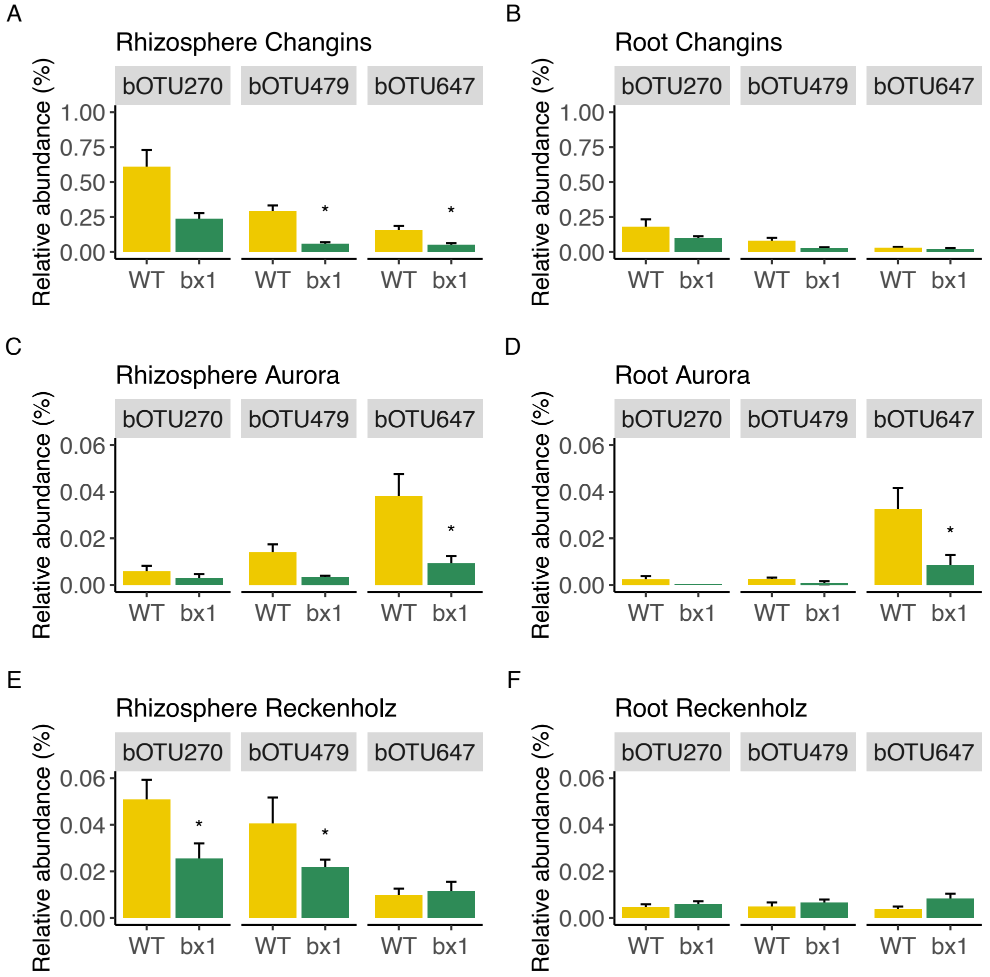


**Figure S10 | Abundance of *Methylophilaceae* bOTUs across compartments and locations**

Bar graphs display the mean abundance (± s.e.m.) of the *Methylophilales* bOTU479 and the two *Methylophilaceae* bOTUs #270 and #647 in rhizosphere (A,C,E) and root (B,D,F) samples of wild-type (WT) and *bx1* mutant lines at all three locations. Asterisks mark significant differences between WT and *bx1* as determined by edgeR analysis (FDR < 0.05, **Table S12**).

### SUPPLEMENTARY TABLES:

**Table S1 | Deposition of raw sequence data**

| Location | Data | MiSeq | Study accession[1] | Sample ID[1] | Publication[2] |
|---|---|---|---|---|---|
| Reckenholz | 16S | Run 09 | PRJEB27162 | SAMEA6521205 | Bodenhausen et al., 2019 |
| Reckenholz | ITS | Run 13 | PRJEB36599 | COMING | this study |
| Aurora | 16S | Run 11 | PRJEB20127 | SAMEA4698767 | Hu et al., 2018 |
| Aurora | ITS | Run 12 | PRJEB36599 | COMING | this study |

[1]At the European Nucleotide Archive (http://www.ebi.ac.uk/ena)
[2]In our laboratory, we typically sequence several different experiments in a single MiSeq run. Therefore, it is possible that the raw data was deposited previously in the context of an earlier study.

**Table S2 | Taxonomic analysis comparing locations and compartments**

The table reports for all sample groups (location * compartment) the F statistic together with the corresponding P value (adjusted for multiple hypothesis testing, following Bonferroni Hochberg) and the relative average abundances [%]. Locations are abbreviated with Ch=Changins, Au=Aurora and Re=Reckenholz. Compartments are abbreviated with So=Soil, Rh=Rhizosphere and Ro=Root. The columns with the posthoc Tukey test results is indicated with "_T" for each sample group. Different letters are used for groups that differ significantly in their abundances (alpha=0.01).

-> See supplementary excel file (Table_S2_Phylum_statistics.xlsx)

**Table S3 | Alpha diversity analysis comparing locations and compartments**

Statistics of Shannon diversity between compartments (soil, rhizosphere and root) at each location (Changins, Aurora and Reckenholz; Tukey HSD test). Different letters correspond to groups that are significantly different from each other in Shannon diversity at $P < 0.05$.

| Location | Compartment | Bacteria | | Fungi | |
|---|---|---|---|---|---|
| Changins | Soil | 1295 | ef | 99.44 | cde |
| | Rhizosphere | 1012 | de | 78.69 | cd |
| | Root | 520.6 | bc | 22.74 | b |
| Aurora | Soil | 1468 | ef | 79.43 | c |
| | Rhizosphere | 687.2 | cd | 104.7 | de |
| | Root | 401.5 | b | 14.2 | a |
| Reckenholz | Soil | 1691 | f | 107.5 | e |
| | Rhizosphere | 241.1 | a | 71.64 | c |
| | Root | 151.2 | a | 19.59 | b |

**Table S4 | Beta diversity analysis comparing locations and compartments**

Effects of location and compartment on beta diversity of bacterial and fungal diversity. R2 and *P*-values of PERMANOVA on Bray-Curtis distances are presented.

| | **location** | | **compartment** | |
|---|---|---|---|---|
| | R2 | *P* | R2 | *P* |
| Bacteria | 0.2325 | **0.001** | 0.1969 | **0.001** |
| Fungi | 0.2178 | **0.001** | 0.2561 | **0.001** |

**Table S5 | Taxonomic analysis comparing BX effects in each compartment**

The table reports for bacteria and fungi and for each compartment the F statistic together with the corresponding P value (adjusted for multiple hypothesis testing, following Bonferroni Hochberg). The table also displays the relative average abundances [%] per sample group (location * genotype). Locations are abbreviated with Ch=Changins, Au=Aurora and Re=Reckenholz. Genotypes are abbreviated with WT=wildtype and bx for the bx1 mutant. The columns with the posthoc Tukey test results is indicated with "_T" for each sample group. Different letters are used for groups that differ significantly in their abundances (alpha=0.01).

-> See supplementary excel file (Table_S5_Phylum_BX_statistics.xlsx)

**Table S6 | Alpha diversity analysis comparing BX effects in each compartment**

Effect of BX production (Wild type vs. mutant) on Shannon diversity within compartments. *F* and *P*-values (if significant, then in bold) from ANOVA are presented.

| | **Root** | | **Rhizosphere** | | **Soil** | |
|---|---|---|---|---|---|---|
| | F | *P* | F | *P* | F | *P* |
| Bacteria | 0.0087 | 0.9219 | 1.882 | 0.1744 | 0.0028 | 0.9579 |
| Fungi | **14.11** | **0.0003** | 0.5741 | 0.451 | 0.1515 | 0.6987 |

**Table S7 | Beta diversity analysis comparing BX and location effects in each compartment**

Location and genotype effects of BX-exudation on beta diversity of bacterial and fungal communities within compartments. R2 and *P*-values of PERMANOVA on Bray-Curtis distance are presented. Significant effects are in bold.

| | | Soil | | Rhizosphere | | Root | |
|---|---|---|---|---|---|---|---|
| | | R2 | *P* | R2 | *P* | R2 | *P* |
| Bacteria | location | **0.3376** | **0.001** | **0.511** | **0.001** | **0.5354** | **0.001** |
| | genotype | 0.01344 | 0.245 | **0.02244** | **0.007** | **0.01642** | **0.044** |
| Fungi | location | **0.2612** | **0.001** | **0.47** | **0.001** | **0.422** | **0.001** |
| | genotype | 0.01472 | 0.263 | **0.02152** | **0.011** | **0.06145** | **0.001** |

**Table S8 | Beta diversity analysis comparing background and genotype effects in each compartment (Reckenholz experiment)**

Factorial (A) PERMANOVA analysis and (B) pairwise PERMANOVA analysis of bacterial and fungal communities at the Reckenholz location testing for effects of compartment (soil, root and rhizosphere), genetic background (B73, W22) and plant genotype (WT, *bx1*). R2 and *P*-values of PERMANOVA on Bray-Curtis distance are presented. For pairwise PERMANOVA, R2 values are reported and significant (FDR < 0.05) effects marked in bold.

| (A) | **Compartment** | | **Background** | | **Genotype** | |
|---|---|---|---|---|---|---|
| | R2 | *P* | R2 | *P* | R2 | *P* |
| Bacteria | **0.3486** | **0.001** | **0.02734** | **0.004** | **0.03431** | **0.002** |
| Fungi | **0.4987** | **0.001** | **0.02524** | **0.005** | **0.04461** | **0.001** |

| (B) | | ***bx1*(b73) - *bx1*(W22)** | ***bx1*(b73) - B73** | ***bx1*(b73) - W22** | ***bx1*(W22) - B73** | ***bx1*(W22) - W22** | **B73 - W22** |
|---|---|---|---|---|---|---|---|
| Bacteria | Root | 0.07705 | 0.0988 | 0.1365 | 0.1266 | 0.08215 | 0.1203 |
| Fungi | Root | 0.1658 | 0.2182 | 0.2131 | **0.3666** | 0.2743 | 0.1203 |
| Bacteria | Rhizo. | 0.07349 | 0.2425 | 0.05906 | **0.3524** | 0.08487 | 0.2326 |
| Fungi | Rhizo. | 0.1096 | 0.1701 | 0.1374 | **0.2402** | 0.1576 | 0.2042 |

**Table S9 | zOTUs differing by background or genotype (Reckenholz experiment)**

The supplementary table lists, for bacteria and fungi of the root and the rhizosphere, all zOTUs differing significantly between backgrounds (B73, W22) or genotypes (WT, *bx1*) based on edgeR analysis. The table lists the zOTU-ID, taxonomy, the mean log abundance in counts per million [logCPM] and log fold change [logFC] between backgrounds or genotypes and the statistic of the likelihood ratio test [LR], its *P*-value [PValue] and its FDR-corrected *P*-value [FDR].

-> See supplementary excel file (Table_S9_Reckenholz_OTU_statistics.xlsx)

**Table S10 | Beta diversity analysis comparing mutant genotypes in each compartment (Aurora experiment)**

Factorial (A) PERMANOVA analysis and (B) pairwise PERMANOVA analysis of bacterial and fungal communities at the Aurora location testing for effects of compartment (soil, root and rhizosphere) and plant genotypes (WT, *bx1*, *bx2* and *bx6*). R2 and *P*-values of PERMANOVA on Bray-Curtis distance are presented. For pairwise PERMANOVA, R2 values are reported and significant (FDR < 0.05) effects marked in bold.

| (A) | **Compartment** | | **Genotype** | |
|---|---|---|---|---|
| | R2 | *P* | R2 | *P* |
| Bacteria | **0.3697** | **0.001** | **0.08198** | **0.002** |
| Fungi | **0.406** | **0.001** | **0.06434** | **0.019** |

| (B) | | ***bx1* – *bx2*** | ***bx1* – *bx6*** | ***bx1* – WT** | ***bx2* – *bx6*** | ***bx2* – WT** | ***bx6* – WT** |
|---|---|---|---|---|---|---|---|
| Bacteria | Root | **0.1405** | **0.1383** | **0.1221** | **0.1289** | **0.196** | **0.1524** |
| Fungi | Root | 0.09296 | 0.1062 | **0.2069** | 0.07716 | 0.1677 | 0.1223 |
| Bacteria | Rhizo. | 0.1355 | 0.2009 | 0.0875 | **0.337** | 0.08852 | 0.2543 |
| Fungi | Rhizo. | **0.1146** | 0.0921 | 0.0875 | 0.0839 | **0.1012** | 0.0896 |

**Table S11 | zOTUs differing by mutant genotypes (Aurora experiment)**

The supplementary table lists, for bacteria and fungi of the root and the rhizosphere, all zOTUs differing significantly between among WT and the mutant lines based on edgeR analysis. The table lists the zOTU-ID, taxonomy, the mean log abundance in counts per million [logCPM] and log fold change [logFC] between WT and *bx1* and the statistic of the likelihood ratio test [LR], its *P*-value [PValue] and its FDR-corrected *P*-value [FDR].

-> See supplementary excel file (Table_S11_Aurora_OTU_statistics.xlsx)

**Table S12 | BX-sensitive zOTUs across all locations**

The supplementary excel table lists for bacteria and fungi of the root and the rhizosphere all zOTUs differing significantly between WT and *bx1* mutant lines for each location based on edgeR analysis. That table lists the location, the zOTU-ID, taxonomy, the mean log abundance in counts per million [logCPM] and log fold change [logFC] between WT and *bx1* and the statistic of the likelihood ratio test [LR], its *P*-value [PValue] and its FDR-corrected *P*-value [FDR].

-> See supplementary excel file (Table_S12_BX_OTU_statistics.xlsx)
